# Life history evolution facilitates trophic diversification

**DOI:** 10.1101/2025.03.25.645227

**Authors:** Catalina Chaparro-Pedraza, Claudia Bank

## Abstract

To what extent is biodiversity shaped by environmental conditions, and to what extent is it the result of self-organization? Both natural processes and structural features may contribute to promoting diversity. Here, we show that one such process, namely natural selection, and a structural feature, namely organismal life history, interact in a feedback mechanism that promotes the emergence of diversity. We illustrate how this mechanism operates using various models of ecological diversification driven by intraspecific resource competition, in which both a niche trait that determines resource use and a life history trait can evolve. We find that natural selection that acts on life history traits leads to increased competition, which, in the presence of ecological opportunity, facilitates niche diversification. As a consequence, the environmental conditions (e.g. productivity) for diversification are more restrictive in the absence of life history evolution than in its presence. Our findings indicate a strong influence of life history evolution on ecological processes that in turn shape the origin of biodiversity. Our results call for a better integration of life history evolution and ecological diversification in theoretical and empirical research.

## Introduction

Ecosystems are complex adaptive systems in which system properties such as species diversity and community structure emerge from interactions among components. Such systems exhibit a large extent of self*-*organization based on a hierarchy of feedback mechanisms in which the outcome of interactions promotes further interactions (*1-4*). While there is a growing awareness that feedback mechanisms are key for the maintenance of biodiversity (*5, 6*), their role in the emergence of biodiversity is unknown.

Most of the diversity of life on Earth is thought to have emerged through ecological diversification (*7, 8*). This process, during which a single ancestral population diversifies into ecologically different species that exploit a variety of niches (*8, 9*), has produced diverse clades through rapid diversification, including Darwin’s finches (*10*), Caribbean anole lizards (*11*), and cichlid fishes (*12*). In these and other examples of ecological diversification, the availability of relatively unexploited ecological niches, known as ecological opportunity, results in a regime of frequency*-*dependent selection that emerges from competition. In such a regime, geographic barriers are not required for diversification when evolution drives a trait to the vicinity of a local fitness minimum, at which selection is divergent and trait diversification occurs (*13, 14*). Competition among individuals is at the core of the ecological diversification process (*15-17*); therefore, any factor modulating the strength of intraspecific competition influences the occurrence of ecological diversification.

Indeed, a variety of factors are known to influence the strength of competition in a population and, thus, the occurrence of ecological diversification. Much research has investigated how favorable environmental conditions, such as those causing high productivity or low mortality, facilitate ecological diversification by strengthening intraspecific competition (*18-21*). However, the possibility that evolutionary processes, such as natural selection, alter intraspecific competition and thereby influence diversification has not been explored. This is because existing diversification models and empirical research have focused on the evolution of traits subjected to divergent selection, e.g. traits associated with niche specialization, whereas other traits received little attention. If natural selection, by driving changes in traits that are not subjected to divergent selection, strengths intraspecific competition, it could facilitate diversification. However, it is unknown whether natural selection tends to favor conditions that promote diversification and, thus, high diversity (*9*).

Key insights into this problem come from previous theoretical work on life history evolution, which showed that the evolution of life history traits can alter intraspecific competition (*22*). Life history traits such as the time organisms take to grow, when they become mature, and how many offspring of a particular size they produce, influence their reproductive success. Natural selection can drive changes in these life history traits, shaping the life cycle of organisms to increase their fitness in the face of ecological challenges posed by the environment (*23-25*). After colonizing a novel habitat, natural selection may drive life history adaptation in a population by replacing life history phenotypes of low reproductive success with phenotypes of higher success. This gradual increase in reproductive success results in stronger intraspecific competition (*22*), potentially facilitating diversification. To date, we do not know how life history evolution, by affecting intraspecific competition, may alter ecological diversification driven by niche specialization.

Here, we identify life history evolution as a general facilitator of ecological diversification by analyzing three fundamentally different theoretical models. Each model considers ecological niches representing ecological opportunity and a niche trait that determines resource use. (Figure 1A). The models differ in the description of the population dynamics, the evolving life history trait (Figure 1B), and other assumptions underlying organismal processes (e.g. a linear functional response describes individual feeding rate in Model 1 and Model 3 versus a saturating functional response in Model 2). We thus demonstrate that the facilitating effect of life history evolution on diversification holds beyond specific model assumptions and life history traits. Our results thus provide a robust quantitative link between life history evolution and trophic diversification.

**Figure 1.**
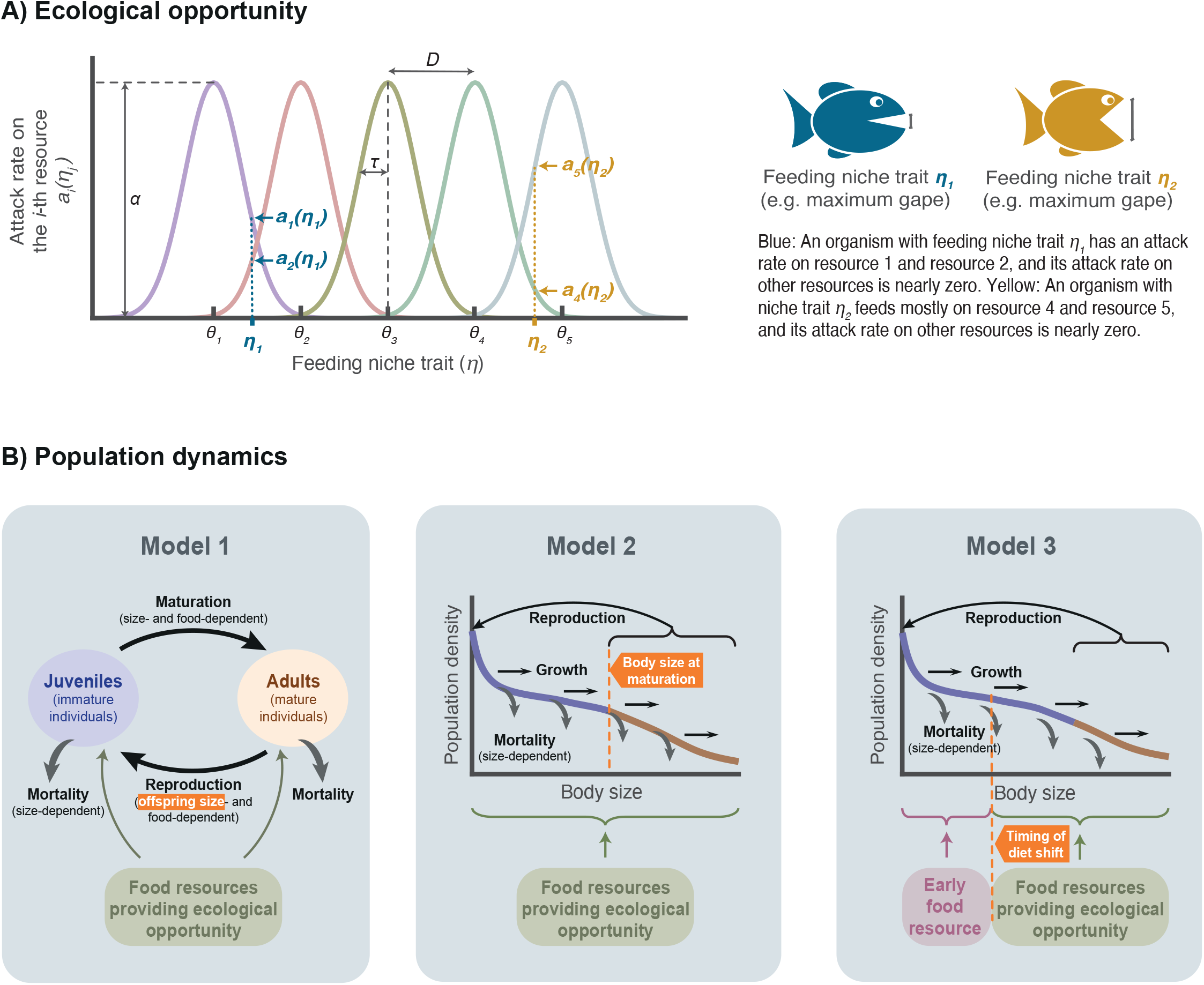
Structure of the models. A) Our models consider the existence of multiple ecological niches, or rather the resources that form the niches, prior to the arrival of an ancestral population. To consume each resource, there exists an optimal feeding niche trait *θ*_*i*_. Organisms differ in the feeding niche trait (e.g., the maximum gape in fish or reptiles, or the bill size in birds, which determines the size of the food particules that they can ingest). B) We study three fundamentally different models. In Model 1, the population is structured by two life stages: juveniles and adults; whereas in Model 2 and Model 3, the population is structured by body size, which changes over an individual’s lifetime. The models also differ in the evolving life history trait, indicated in the orange boxes. In Model 3, competition for food and access to ecological opportunity occurs only after the timing of the diet shift (organisms do not compete for the early food resource).

## Models and results

We formulate and analyze three alternative models to examine how natural selection on life history traits influences ecological diversification. The core of each model is the description of various organismal processes, that is, feeding, growth, reproduction, and mortality, as a function of the environment (food availability) and the organism itself. The ecological dynamics emerging from this description determine the local fitness landscape and, thus, the evolution of life history and feeding niche. Applying adaptive dynamics techniques (*26, 27*), we identify the conditions that enable diversification (analytically derived for Model 1) and simulate the evolutionary trajectories of diversifying lineages.

### Ecological opportunity and food resource use in the three models

Building on existing theory on ecological diversification (*18, 28-30*), our models consider *n* food resources with density *F*_*i*_ (*i* = 1, …, *n*), and *m* emerging consumer ecomorphs (*j* = 1, …, *m*), each with life history trait *l*_*j*_ and feeding niche trait η_*j*_ determining their resource use. In the absence of consumers, resource density dynamics follow *dF*_*i*_/*dt* = *ρ*(*F*_*i max*_ − *F*_*i*_), where *ρ* and *F*_*i max*_ are the renewal rate and the carrying capacity of the *i*th resource, respectively. The total productivity is the sum of the product of the carrying capacities and the supply rate of the resources, such that *P* = *ρ* ∑_*i*_ *F*_*i max*_.

Assuming that there exists an optimal trait value *θ*_*i*_ to consume each resource, these optimal traits are ordered along a one*-*dimensional niche trait space (i.e. *θ*_1_ < *θ*_2_ < ⋯< *θ*_*n*_), separated by a distance *D*. The attack rate, *a*_*i*_(η_*j*_), of an ecomorph with feeding niche trait η_*j*_ on the *i*th resource equals the maximum attack rate *α* when its feeding niche trait η_*j*_ equals *θ*_*i*_, and decreases in a Gaussian manner as *η*_*j*_ moves away from *θ*, that is *a*_*i*_(*η*_*j*_) = *α* exp [−(*η*_*j*_ − *θ*_*i*_) _2_ / (_2_τ) _2_] (figure 1A). In this expression, τ determines the width of the Gaussian function, and hence, when it is small, an ecomorph must be highly specialized to successfully attack resource *i*. Consequently, we assume a tradeoff between feeding on alternative food resources, in which the specialization on one food resource is at the expense of specialization on the others (*31*). Such tradeoffs have been generally observed in heterotrophic organisms, including bacteria (*32*), insects (*33*), and vertebrates (*34, 35*).

### Model 1: Diversification and evolution of offspring size

Model 1 considers an ecomorph population *j* that is composed of juvenile and adult individuals. Individuals are born with size *l*_*j*_ and mature (i.e., enter the adult stage) when they reach size *w*. The feeding rate of juveniles of the *j*th ecomorph is 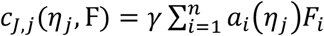 and of adults is 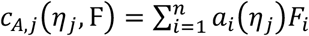, with F = (*F*_1_ … *F*_*n*_). The factor *γ* scales the foraging capacity of juveniles such that, when *γ* > 1 (resp. γ < 1), juveniles feed at a faster (resp. slower) rate than adults. The flux of energy that juveniles allocate to somatic growth is *φ*_*j*_(*η*_*j*_, F) = max(ε *c*_*J,j*_(*η*_*j*_, F) − *ν*, _0_), where *ε* is the assimilation efficiency and *ν* is the metabolic cost. Hence, somatic growth only increases when the assimilated food exceeds the metabolic cost *ν*. De Roos and Persson showed that the maturation rate depends on the somatic growth, the mortality rate, and the size range over which an individual grows in the juvenile stage (see Box 3.1 in ref (*36*)), as given by:

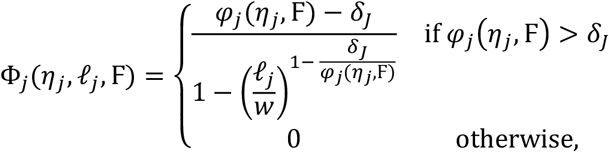

where *δ*_*J*_ is the juvenile mortality rate. The maturation rate thus decreases as the range over which an individual grows in the juvenile stage increases; that is, the ratio *𝓁*_*j*_/w decreases. Adults only reproduce when assimilated food exceeds the metabolic cost incurred by adults, *ν*, such that fecundity is β_*j*_(*η*_*j*_, *𝓁*_*j*_, F) = max(s*c*_*A,j*_(*η*_*j*_, F) − *ν*, _0_)/*𝓁*_*j*_. Fecundity is divided by the offspring size, reflecting the well*-* known tradeoff between offspring number and size that has been ubiquitously observed across diverse plant and animal taxa (*23, 37-39*). Additionally, juvenile survival was documented to increase with offspring size in wild populations of diverse animal species (*40*) because larger organisms often have greater survival than smaller conspecifics (*41-52*). Thus, we assume that juvenile mortality decreases with offspring size, following *δ*_*J*_(*𝓁*_*j*_) = *δ*_max_exp(−*𝓁*_*j*_), where *δ*_max_ is a boundary of maximum possible mortality. Adults die at a rate *δ*_*A*_. Starvation mortality is not considered because, at the ecological equilibrium, a population is viable (i.e., its density is positive) only if starvation conditions do not occur, i.e., if s*c*_*J,j*_ > *ν* and s*c*_*A,j*_ > *ν*.

Based on the assumptions described above, the dynamics of juveniles *J*_*j*_ and adults *A*_*j*_ of the *j*th ecomorph, as well as of food resources, follow

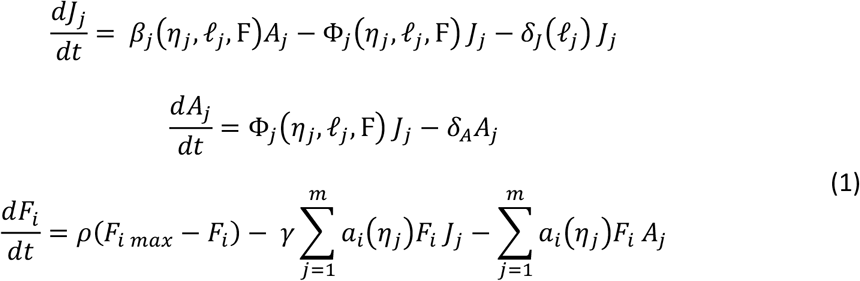

The total population density, *N*_*j*_ = *A*_*j*_ + *J*_*j*_, varies following

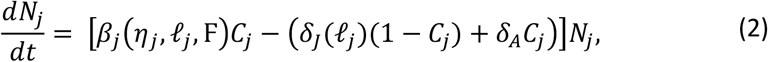

where *C*_*j*_ is the fraction of adults (for the ecological dynamics rewritten in terms of *N*_*j*_ and *C*_*j*_, see SI1: eq. SI1.1). The per capita growth rate, and thus the fitness of the *j*th ecomorph, is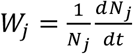. Using this fitness expression and following adaptive dynamics techniques (*26, 27*), we obtained analytical expressions for the selection gradient and the curvature of the fitness landscape when both the feeding niche trait and the offspring size evolve (see SI1: eq. SI1.4*-*SI1.6). For a population encountering an environment with two different food resources with carrying capacities *F*_1 *max*_ = *F*_2 *max*_, we obtained analytical expressions for the food and population densities as well as the fraction of adults at the ecological equilibrium (see SI2). Using these quantities and the expressions for the selection gradient and the curvature of the fitness landscape, we analytically derived the conditions that enable diversification (see SI3) and determined their relationship with the life history trait (see SI4).

Additionally, we simulated the evolutionary trajectories of a diversifying lineage that encounters an environment with ecological opportunity represented by two (figure 2) or more (figure 3) different food resources. To do so, we used eq. SI1.1*-* SI1.3 and parameter values in Table SI5.1 (see model parameterization in SI5). Moreover, we evaluated whether a diversification event occurs by calculating the curvature of the fitness landscape (using eq. SI1.6) when directional selection ceases (i.e., when the change in the feeding niche trait is smaller than 1E*-*8 per evolutionary time step). If the curvature of the fitness landscape indicated that the population experiences disruptive selection in the feeding niche trait (selection in the life history trait is always directional or stabilizing), population *j* was split into two distinct ecomorph populations with trait values 0.1% larger and smaller than *η*_*j*_, and abundance equal to *N*_*j*_/_2_. Subsequently, the evolutionary trajectories were simulated simultaneously for the two ecomorph populations until directional selection in the feeding niche trait ceased, and the curvature of the local fitness landscape was evaluated for each ecomorph. Each simulation finished when all ecomorph populations experienced stabilizing selection. Throughout each simulation, we followed the number of ecomorphs, their traits, and their densities, as well as the densities of resources, which we used to calculate the fitness landscape.

**Figure 2.**
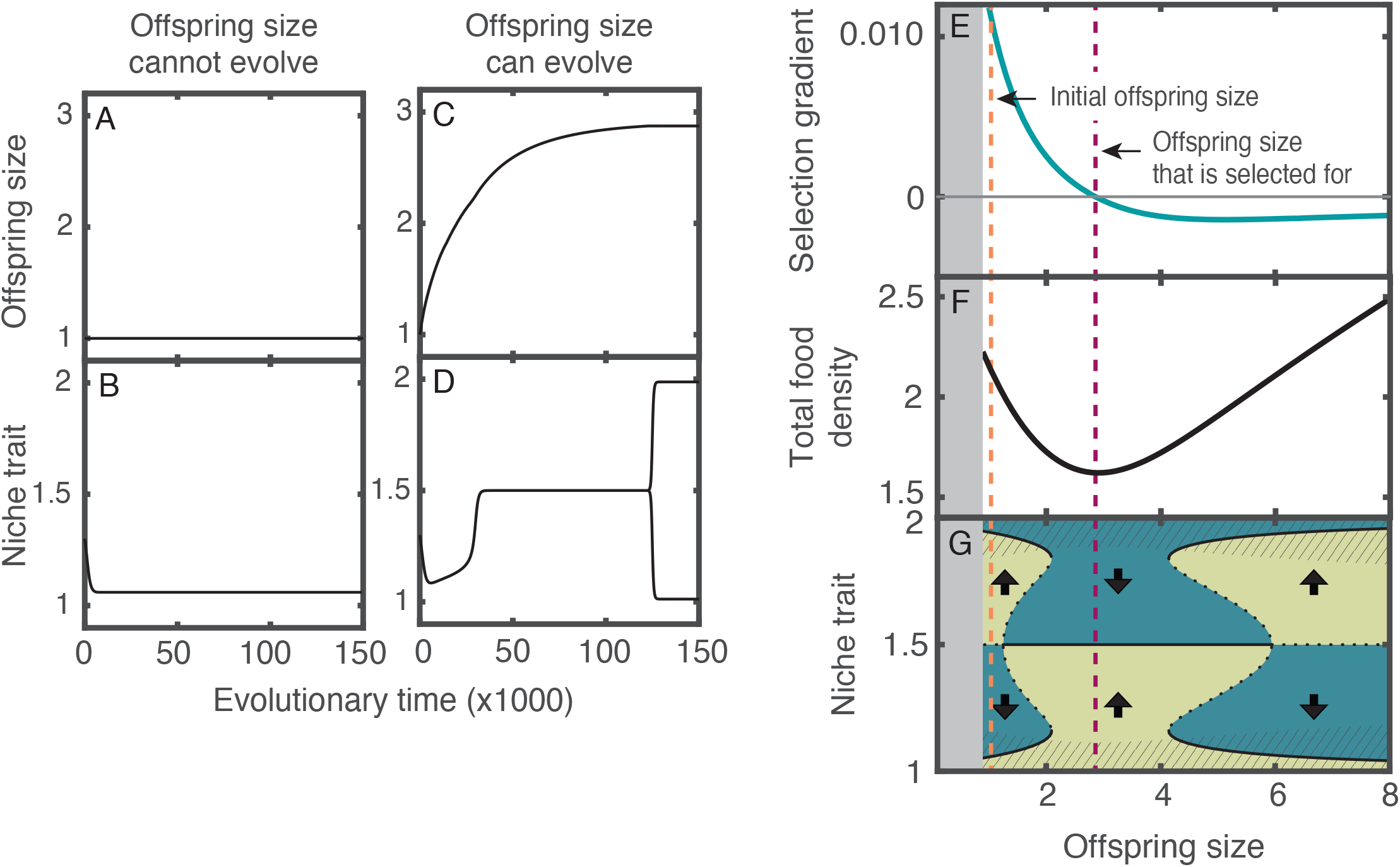
The evolution of offspring size enables diversification. Evolutionary dynamics of a population that colonizes an environment with two food resources (optimal niche traits to feed on resources are *θ*_*1*_ *= 1, θ*_*2*_ *= 2*) A, B) when the offspring size cannot evolve, and C, D) when it can evolve. E) Selection gradient with respect to the offspring size of the colonizing population. F) Sum of the food densities at the ecological equilibrium as a function of the offspring size. G) Fitness landscape of the niche trait as a function of the offspring size. Diversification occurs only when the offspring size of the population is near the optimal trait (indicated with purple vertical dashed line) because the niche trait evolves towards a fitness minimum, where the trait equals 1.5 (arrows indicate the direction of selection). Evolutionary equilibriums (black lines; solid when stable and dotted when unstable) in the striped regions are fitness maxima, whereas outside these regions they are fitness minima. In the grey region in the left, the population is extinct. In A-D, the initial offspring size and niche trait are 1 and 1.3, respectively. In F, the niche trait value is fixed and equal to 1.4. However, regardless of the niche trait value, the food density is always depleted to the lowest level at the value of the offspring size that is selected for (analytical proof in SI4). Parameter values as in table SI5.1.

**Figure 3.**
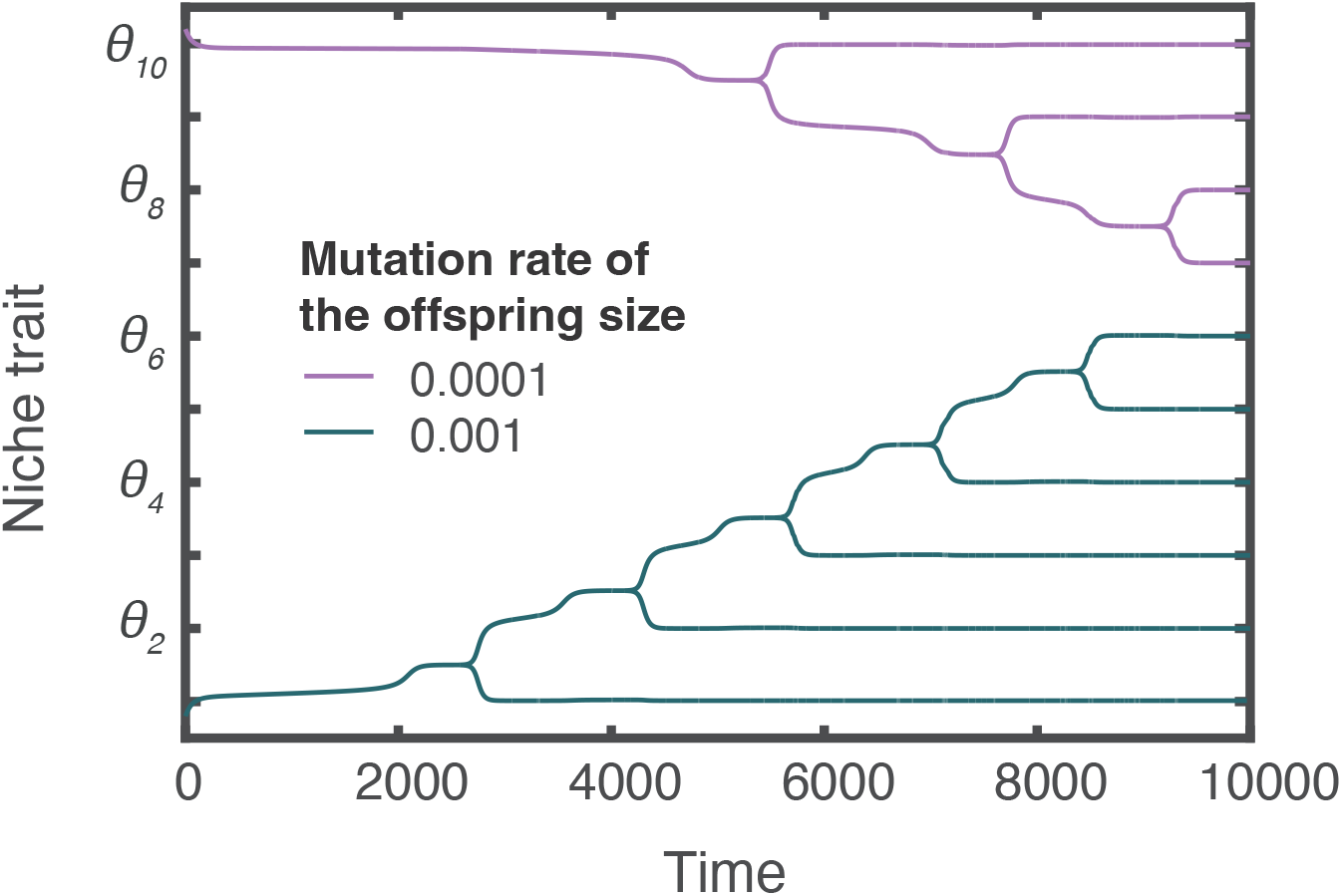
The speed of offspring size evolution influences lineage diversity. Evolutionary trajectories of two lineages colonizing an environment with 10 different unexploited food resources. The lineage with a higher mutation rate in offspring size results in 1.5-fold higher diversity than the lineage with lower mutation rate (the former diversifies into six different ecomorph populations, whereas the latter diversifies into four). The total productivity is 2 gL^-1^ unit time^-1^. The feeding trait values of the ancestral populations are 0.8 and 10.2. The offspring size of both ancestral populations is 1. Parameter values as in table SI5.1.

### By altering the stability of evolutionary equilibria, life history influences the conditions for diversification

For diversification to occur, theory proposes that a population must experience disruptive selection for a significant amount of time (*13*). This is possible when natural selection drives a population towards a local minimum of the fitness landscape, causing intermediate phenotypes to have a fitness disadvantage compared with more extreme phenotypes (*14, 53*). Two conditions need to be satisfied for diversification to occur in a population colonizing an environment with two different food resources:

<u>Condition 1</u> (condition for mutual invasibility according to Geritz et al. (*53*)): The mean phenotype of the population at an evolutionary equilibrium must be a local fitness minimum. A population satisfies this condition when:

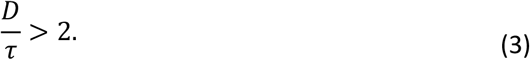

(see analytical derivation in SI3: eq. SI3.19). In our model, the ratio *D*/τ determines the strength of the tradeoff between feeding on the alternative food resources. Eq. 3 thus implies that this tradeoff needs to be sufficiently strong to induce disruptive selection (*28, 54*).

<u>Condition 2</u> (condition for convergence stability according to Geritz et al.(*53*)): The evolutionary equilibrium at which the population experiences disruptive selection must be an attractor of the evolutionary dynamics, or in other words, natural selection must be able to drive a population toward this evolutionary equilibrium. This condition is satisfied when productivity exceeds a given threshold:

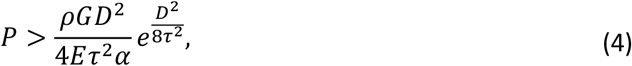

where *E* = ε *C*/*𝓁* is a scaling factor of the population birth rate and *G* = *νC*/*𝓁*+ *δ*_*A*_*C* + *δ*_*J*_(*𝓁*)(1 − *C*) is the rate of biomass loss through various processes (e.g. metabolic maintenance, mortality). These two compound variables regulate population growth, as shown by rewriting eq. 2 as 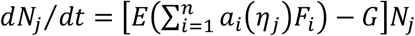. Eq. 4 implies that there exists a minimum productivity of the environment that enables the evolutionary equilibrium at which the population experiences disruptive selection to be an attractor of the evolutionary dynamics (see analytical derivation in SI3: eq. SI3.21). This threshold of minimum productivity depends on simple parameters, *D*, τ, *ρ* and *α*, and the compound variables, *E* and *G*. Large values of *E* or small values of *G* reduce this threshold, enabling diversification over a larger range of productivities. This is because by increasing *E*, the population birth rate increases. Analogously, by reducing *G*, the biomass loss decreases. Consequently, in both cases, the population density increases, leading to stronger intraspecific competition that, in turn, facilitates diversification.

Life*-*history evolution through offspring size in Model 1 does not affect Condition 1, but it affects Condition 2. Specifically, evolution of offspring size alters the stability condition that determines when selection drives the feeding niche trait to the value where it becomes disruptive. This is due to the effect of offspring size evolution on the population birth rate through *E* and the biomass losses through *G*. These effects are direct; for instance, the tradeoff between number and offspring size directly affects the birth rate, and the size*-*dependent mortality experienced by the offspring impacts the biomass losses. In addition, these effects are indirect, mediated by changes in population composition, *C*. Hence, by influencing the population composition, its birth rate, and its losses, the offspring size determines whether the feeding niche trait value at which selection is disruptive is an attractor or repeller of the evolutionary dynamics and thus dictates whether diversification occurs or not.

### The evolution of life history traits enables diversification

To understand how life history evolution influences ecological diversification, we simulated two scenarios: 1) a null model in which only the niche trait can evolve (i.e. the mutation rate for offspring size is set to zero), and 2) an alternative model in which both traits, niche and offspring size, can evolve. In the first scenario, in which offspring size remains constant (figure 2A), the niche trait of the ancestral population evolves towards one of the resource feeding optima (figure 2B). At this point, diversification cannot occur because selection is stabilizing. In the second scenario, the niche trait is initially driven toward one of the resource feeding optima (between time 0 and 5000 in the simulation); however, later, when the offspring size approaches an intermediate trait value (offspring size of 1.5 in figure 2C), the direction of selection changes for the niche trait. As a result, the niche trait evolves to the value between the two optima. At this point, selection becomes disruptive; therefore, the population undergoes a diversification event in the niche trait. After diversification, the niche traits of the two resulting ecomorph populations diverge. Each population then evolves towards the nearest optimal trait value to feed on the resources (figure 2D).

To better understand the mechanism by which life history evolution enables diversification, we analyzed the fitness landscape of the niche trait experienced by the colonizing population. Our analysis revealed that ecological diversification can only occur when the offspring size is near its optimal value (purple vertical line in figure 2E). In the neighborhood of the optimal offspring size, food resources are strongly depleted, causing strong intraspecific competition (figure 2F). As a consequence, the niche trait value between the two optima to feed on the resources (niche trait 0.5 in figure 2) is an attractor of the evolutionary dynamics, and thus, the niche trait evolves towards this attractor (when offspring size is between 1.2 and 5.9 in figure 2G). Because this niche trait value is a local fitness minimum, selection becomes disruptive in its vicinity, leading to diversification. Conversely, when the offspring size is far from its optimal value, this niche trait value is a repeller of the evolutionary dynamics, and consequently, evolution drives the niche trait away from it (offspring size below 1.2 and above 5.9 in figure 2G).

In line with the conditions for diversification presented in the previous section, the fitness landscape analysis revealed that the niche trait value corresponding to a local fitness minimum must be an attractor of the evolutionary dynamics for diversification to occur. This is possible when the trait value on which a local fitness minimum resides is either 1) a global attractor of the evolutionary dynamics, and thus, evolution drives the niche trait towards it regardless of the trait value of the initial population (e.g., when offspring size is larger than 2.2. and smaller than 4.2 in figure 2G), or 2) a local attractor, and thus, evolution can only drive the niche trait towards it if the trait value of the initial population is in its neighborhood (e.g., when offspring size is between 1.2 and 2.2 or between 4.2 and 5.9 in figure 2G).

### The evolutionary rate of life history traits influences diversification and the ensuing diversity

The effect of life history traits on the fitness landscape of the feeding niche trait suggests that the speed at which life history traits evolve affects diversification. To test this, we simulated the evolutionary trajectories of two lineages colonizing an environment with ten different unexploited niches. These lineages differ in the mutation rate of the life history trait (i.e. the offspring size) and thus the rate of change of this trait, but have equal mutation rates of the feeding niche and access to the niches (both lineages are seeded at the beginning of the simulation in the extremes of the trait space). The simulation shows that the diversification process initiates earlier in the lineage with the highest mutation rate (figure 3). This is because, in this lineage, the rate of change of the offspring size is higher, enabling natural selection to drive this trait towards the optimal value faster. Then, in the neighborhood of the optimal offspring size, natural selection drives the feeding niche trait towards the value in between two optima (e.g., between *θ*_1_ and *θ*_2_ at time 2500), where selection becomes disruptive, enabling diversification. Conversely, in the lineage with the lower mutation rate, the evolutionary process driving changes in the offspring size is slower. Consequently, it takes twice as long for the feeding niche trait to reach the value in between two optima (e.g., between *θ*_9_ and *θ*_10_ at time 5000). As a result of the earlier diversification process, the lineage with the highest evolutionary rate in the life history trait occupies more niches, resulting in a higher diversity.

### Local adaptation facilitates diversification

We next explored how variation in environmental conditions across habitats affects life history evolution and diversification. To this end, we simulated alternative environments that differ in the mortality risk experienced by juveniles. Because juvenile mortality decreases as the offspring size increases, in more risky environments natural selection favors a larger offspring size. The optimal offspring size is therefore shifted towards a larger trait value when the mortality risk of juveniles is higher (top row in figure 4A). Interestingly, the range of productivity levels over which diversification can occur is larger when the offspring size is near its optimum than when it is far from the optimum (bottom row figure 4A). Hence, by driving offspring size towards the optimal trait value, natural selection reduces the threshold of minimum productivity required for diversification to occur, facilitating diversification. We prove this effect analytically in SI4 by showing that eq. 4 always assumes its minimum value when the offspring size is at its optimum.

**Figure 4.**
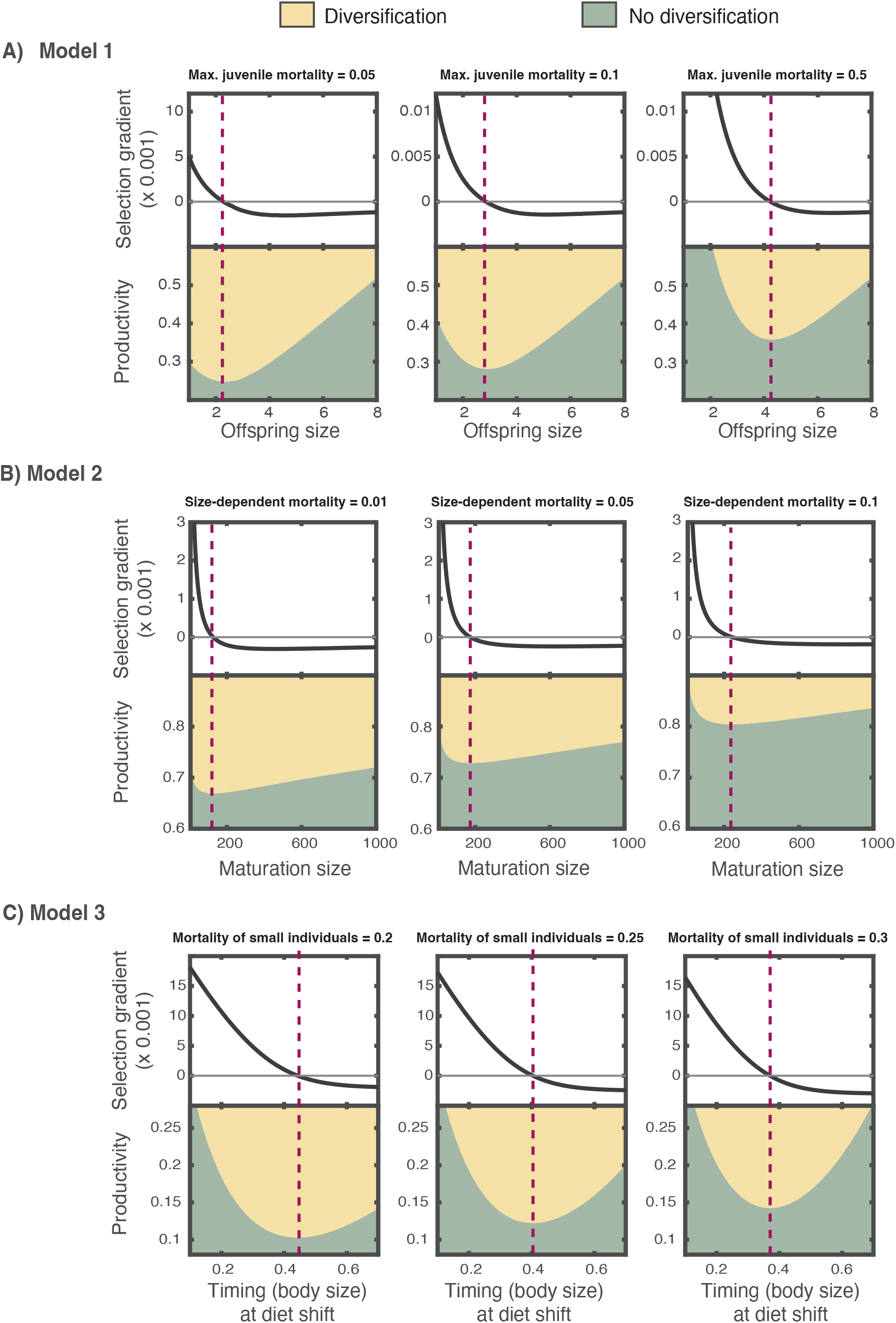
Local adaptation in life history traits facilitates diversification. The range of productivities over which diversification is enabled increases in the neighborhood of the optimal life history trait (i.e. where the selection gradient of the offspring size in A, maturation size in B, and timing at diet shift in C equals zero). For each model, three scenarios that differ in the mortality risk experienced by juveniles in A, all individuals in B or the small individuals in C are simulated. Other parameter values of model 1 as in table SI5.1, of model 2 as in table SI6.1 and of model 3 as in table SI6.2.

### Model 2 and 3: Diversification and evolution of maturation size and timing of diet shif

To test the generality of our results, we examined how natural selection influences ecological diversification by driving changes in other life history traits. In model 2, we study the effect of the evolution of body size at maturation, which determines the transition of energy allocation from growth to reproduction (figure S1). Here, the optimal energy allocation strategy, and thus the optimal size at maturation, depends on the tradeoff between reproductive investment and size*-*dependent survival (*25*). Using this model, we simulated alternative environments that differ in the magnitude of size*-*dependent survival to understand how environmental variation in survival affects the evolution of body size at maturation and diversification. In Model 3, we study the consequences of the evolution of the timing of a diet shift. This trait is associated with a tradeoff between growth potential and survival (*55*), and is thus fundamental in determining individual fitness (*56-59*). Using this model, we simulated alternative environments that differ in the mortality experienced by small individuals (i.e. individuals feeding on the early food resource) to examine how environmental variation in mortality in early life influences the evolution of the timing of a diet shift and diversification. For both models, we leveraged existing ecological models (*35, 60*) that we extended to include evolution of the life history and feeding niche trait. See SI6 for a detailed description of the two models. All analyses were performed using the PSPManalysis software package (*61*), which allows for the evolutionary analysis of population models structured by body size (*62, 63*).

### General patterns: optimal life histories maximize the potential for diversification

Across all studied models, we found that the productivity necessary for diversification increases with the distance of the life history trait from its optimum (figure 4). Our analysis of the two alternative model systems thus generalizes our previous findings from Model 1. The three models greatly differ in their formulation of the population dynamics, the evolving life history traits, and their assumptions underlying life history processes. Despite these differences, all models similarly suggest that natural selection, by driving local adaptation in life history, facilitates trophic diversification.

## Discussion

Here, we showed that life history adaptation results in stronger intraspecific competition for food resources, which in turn promotes diversification. Both theory (*13, 14*) and empirical evidence(*15-17*) demonstrated that by causing negative frequency*-*dependent interactions, intraspecific resource competition can be a source of disruptive selection. This may increase phenotypic diversity within a population (*64*) or result in ecologically driven sympatric speciation in case barriers to gene flow evolve between divergent phenotypes (*65, 66*). In either case, diversity is enhanced. Previous studies hypothesized that natural selection, by operating at the population level, may have consequences at the community level that ultimately promote diversity (*67-69*). However, the underlying mechanisms remained unclear. Our results show that, after colonization of a new environment, life history evolution can facilitate trophic diversification, promoting diversity. Diversity, in turn, was shown to influence dispersal, and thus colonization rates, by altering ecological interactions that exert selection on dispersal*-*related traits (*70-75*). A self*-*reinforcing feedback loop promoting diversity emerges in case the outcome of the altered interactions promotes dispersal (figure 5). Our work suggests that natural selection operating on life history traits is at the core of this feedback, emerging as a driver of self*-*organization in diverse ecological communities.

**Figure 5.**
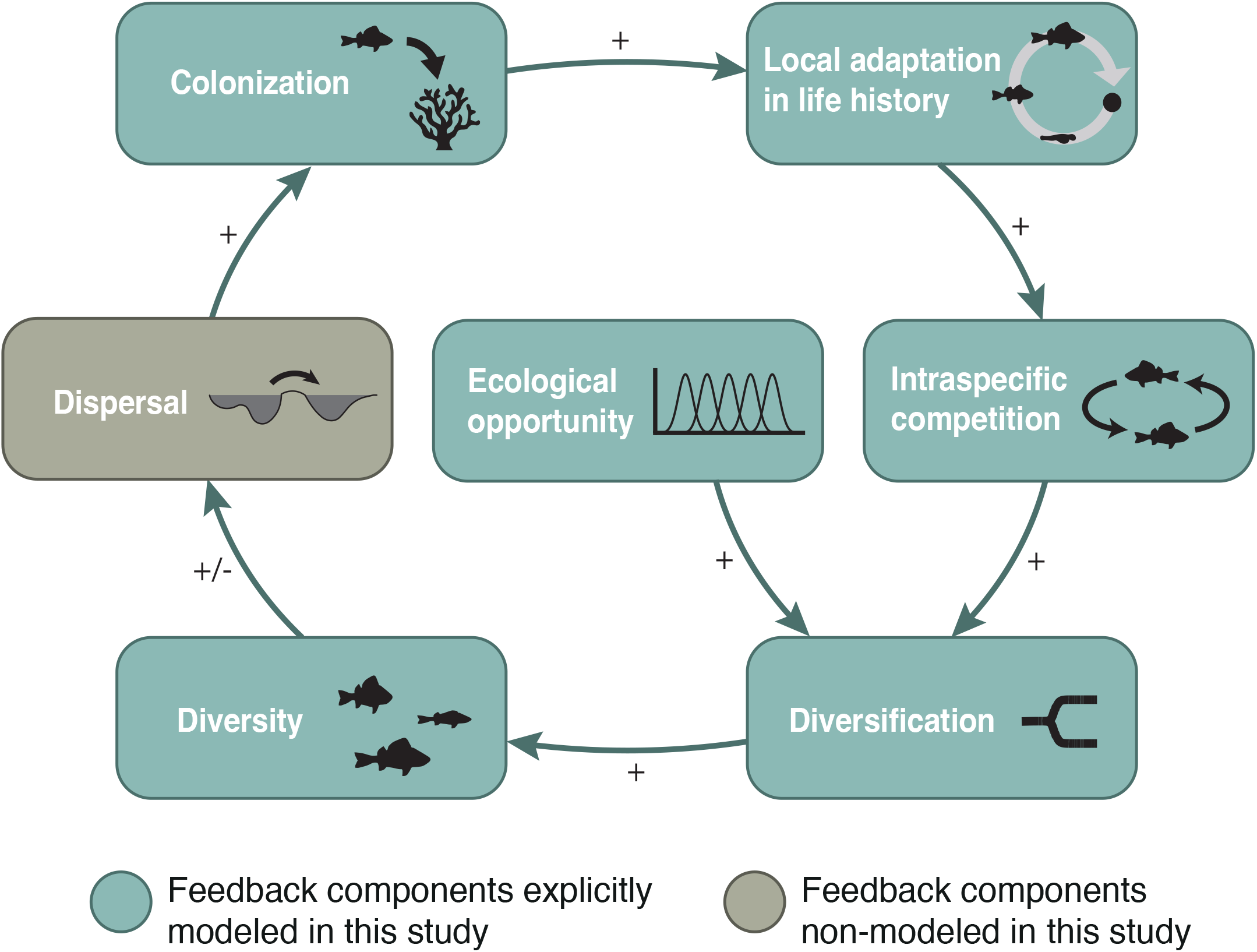
An eco-evolutionary feedback loop promoting diversity. After colonization of a novel environment, local adaptation in life history results in stronger intraspecific competition, which, in the presence of ecological opportunity, induces disruptive selection. As a consequence, ecological diversification driven by niche specialization occurs, increasing phenotypic diversity within populations or species diversity in case barriers to gene flow evolve. In either case, functional diversity is enhanced, which, in turn, can positively influence dispersal and thereby the rate of colonization of novel environments.

We find that a fast rate of adaptation of life history traits accelerates diversification, boosting lineage diversity. Empirical evidence from ray*-*finned fishes, the largest vertebrate clade, may support the validity of this prediction. In this clade, species richness across lineages and diversification rates correlate with the evolutionary rate of body size (*76*). This morphological trait influences multiple life history traits (*77-80*), but it also plays a role in providing access to certain ecological niches. For example, the emergence of piscivorous species in various fish radiations entails an increase in body size (*81-83*). Therefore, disentangling the effects of body size evolution on life history from its effects on ecological traits associated with niche specialization is challenging. Future research is needed to evaluate our prediction, e.g., by studying the evolutionary rates of life history traits, e.g. fecundity*-*at*-*age, maturation size, offspring size, etc., rather than body size, in related radiating and non*-*radiating taxa or taxonomic groups.

Our findings suggest that rapid local adaptation in life history traits facilitates diversification. Anecdotal evidence from Salmonids supports this prediction. Various lineages of these fish have independently diversified producing an array of species exploiting diverse resources within temperate lakes when new habitats have become available (e.g., after the last glaciation) (*83-85*). The extent of local adaptation in salmonids is remarkably high and manifests quickly following the invasion of a new habitat (*86*). Additionally, life history traits in salmonids, such as offspring size, are documented to evolve rapidly in response to a shift in selection pressures (e.g., in the transition from wild to captivity, salmon populations quickly evolve small egg size due to the removal of selection imposed by predation) (*87*). Our results provide an explanation linking fast life history evolution, rapid local adaptation, and propensity to diversify.

Here, we studied ecological diversification leading to specialization to exploit a variety of resources. Classical theory, in contrast, modeled ecological diversification assuming a single resource (e.g., ref (*88-90*)). This model feature makes it impossible for natural selection to drive a trait away from an evolutionary equilibrium at which selection turns disruptive. In other words, one of the conditions for diversification, more precisely, the condition for convergence stability (*53*), is always satisfied in those models. We show that life history traits affect diversification only via this condition. Therefore, by assuming a single food resource, classical models of ecological diversification preclude the possibility that the evolution of life history traits influences diversification. This assumption dominates existing theory on ecological diversification, despite empirical evidence showing that ecological diversification typically involves specialization to exploit diverse resources (*11, 91*). Only a few earlier models have considered multiple resources (*28, 66*), and found that the condition for convergence stability is not always satisfied, as in our model. However, we are not aware of any models that have included evolution of life history traits.

Our findings suggest that lineages with a higher rate of evolution of life history traits are expected to have higher diversification rates. We found this by simulating the evolutionary dynamics of lineages that differ in the mutation rate associated with the life history trait. However, other genetic processes may affect the rate at which life history adaptation occurs in response to natural selection. For instance, gene duplication, recombination, and epigenetic modification can also contribute to creating phenotypic variation that is both heritable and adaptive (*92*). Therefore, by altering the adaptation rate of life history traits, these processes could have a similar effect on diversification as the mutation rate. Future research should consider explicit trait genetic architecture, using, e.g., individual*-*based model simulations (*93*), to investigate the role of these processes in life history evolution and its consequences for diversification.

Central to the mechanism promoting diversity introduced here is the feedback between the individual and its environment: the individuals’ diet collectively affects the food resource availability, which in turn changes the profitability of an individual’s diet. This feedback enables the emergence of a frequency*-*dependent selection regime, which is fundamental to inducing disruptive selection and thus diversification (*13, 14*). In this study, we examined the effect of life history evolution on diversification when the feedback between the individual and its environment results in the depletion of the food resources that provide ecological opportunity. Under this scenario, we found life history evolution to facilitate diversification because the optimal life history strategy coincides with the strategy causing the strongest intraspecific competition for these resources. However, the utilization of additional resources may complicate diversification dynamics. This scenario can occur when only a part of the population has access to ecological opportunity, while the other part is limited by a different food resource. For example, in many pairs of sympatric species of fish (*94*) and anurans (*95*) with complex life cycles, only large individuals are specialized to feed on specific resources, whereas small individuals of multiple species share the same ecological niche. In such cases, if resource competition is stronger among small than among large individuals, small individuals grow slowly, resulting in low recruitment into the life stage that has access to ecological opportunity. As a consequence, competition for the resources that provide ecological opportunity is weak, hindering diversification (*94*). Given that life history adaptation in this scenario is driven by the availability of the resources that provide ecological opportunity as well as another resource, the optimal life history strategy would not necessarily coincide with the strategy causing the strongest intraspecific competition for the resources providing ecological opportunity. Future research will be needed to unveil the consequences of life history evolution for the diversification of organisms with complex life cycles with intraspecific competition for food in multiple life stages.

Uncovering the effect of life history evolution on ecological diversification required the integration of ecology and two bodies of evolutionary theory that, to our knowledge, have been studied separately thus far: life history theory and diversification theory. Life history traits directly affect an organism’s fitness by influencing survival and reproduction. Consequently, their evolution alters population demography, which impacts major ecological processes, such as intraspecific competition, as our study shows. Given that intraspecific competition plays a prominent role in several ecological and evolutionary processes, such as species coexistence (*96*), future research should address the consequences of life history evolution beyond ecological diversification. Understanding these consequences will require the integration of life history theory and other ecological and evolutionary theories. We have demonstrated how such integration unveils a general influence of life history evolution on diversification. Our study highlights the need to synthesize diverse research bodies to study the mechanisms underlying the origin and maintenance of biodiversity.

## References

1. S. A. Levin, Ecosystems. 1, 431–436 (1998).

2. S. A. Kauffman, The origins of order: Self-organiza8on and selecyion in evolution (Oxford University Press, USA, 1993).

3. K. Sneppen, P. Bak, H. Flyvbjerg, M. H. Jensen, Proceedings of the National Academy of Sciences of the United States of America. 92, 5209–5213 (1995).

4. S. E. Jørgensen, H. Mejer, S. Nors, 111, 261–268 (1998).

5. M. P. Veldhuis, M. P. Berg, M. Loreau, H. Olff, Ecological Monographs. 88, 304–319 (2018).

6. D. A. Perry, Trends in Ecology & Evolution. 10, 241–244 (1995).

7. G. Simpson, The Major Features of Evolution (Columbia University Press, New York, 1953).

8. D. Schluter, The ecology of adaptive radiation (OUP Oxford, 2000).

9. R. De-Kayne et al., Cold Spring Harbor Perspectives in Biology, a041448 (2024).

10. P. R. Grant, Ecology and Evolution of Darwin’s Finches (Princeton University Press, 2017).

11. J. B. Losos, S. Gavrilets, Science. 323, 732–738 (2009).

12. W. Salzburger, B. Van Bocxlaer, A. S. Cohen, Annual Review of Ecology, Evolu8on, and Systematics. 45, 519–545 (2014).

13. C. Rueffler, T. J. M. Van Dooren, O. Leimar, P. A. Abrams, Trends in Ecology and Evolution. 21, 238–245 (2006).

14. M. Doebeli, Adaptive diversification (2011).

15. D. I. Bolnick, Evolution. 58, 608–618 (2004).

16. R. Svanbäck, D. I. Bolnick, Proceedings of the Royal Society B: Biological Sciences. 274, 839–844 (2007).

17. E. J. A. Minter, P. C. Wais, C. D. Lowe, M. A. Brockhurst, Biology LePers. 11 (2015), doi:10.1098/rsbl.2015.0192.

18. P. C. Chaparro-Pedraza, G. Roth, O. Seehausen, Ecology LePers. 25, 802–813 (2022).

19. R. Benmayor, A. Buckling, M. B. Bonsall, M. A. Brockhurst, D. J. Hodgson, Evolution. 62, 467–477 (2008).

20. R. Kassen, Annals of the New York Academy of Sciences. 1168, 3–22 (2009).

21. A. R. Hall, N. Colegrave, Proceedings of the Royal Society B: Biological Sciences. 274, 73–78 (2007).

22. S. D. Mylius, O. Diekmann, Oikos. 74, 218–224 (1995).

23. S. C. Stearns, The evolution of life histories (Oxford university press, 1998).

24. J. Torres Dowdall et al., Funct Ecol. 26, 616–627 (2012).

25. D. Roff, Evolution of life histories: theory and analysis (Springer Science & Business Media, 1993).

26. U. Dieckmann, R. Law, Journal of Mathematical Biology. 34, 579–612 (1996).

27. O. Leimar, Evolutionary Ecology Research. 11, 191–208 (2009).

28. C. Rueffler, T. J. M. Van Dooren, J. A. J. Metz, American Naturalist. 167, 81–93 (2006).

29. C. Chaparro-Pedraza, G. Roth, C. Melian, Scientific Reports. 14 (2024), doi:10.1101/2024.03.06.583698.

30. P. C. Chaparro-Pedraza, The American Naturalist. 204 (2024), doi:10.1086/730446.

31. R. Levins, Evolution in changing environments: some theoretical explorations (Princeton University Press, 1968).

32. D. M. Ekkers et al., Molecular Biology and Evolution. 39, 974–984 (2022).

33. R. L. Ware, M. E. N. Majerus, From biological control to invasion: The ladybird Harmonia axyridis as a model species, 169–188 (2008).

34. A. Herrel, J. Podos, B. Vanhooydonck, A. P. Hendry, Functional Ecology. 23, 119–125 (2009).

35. M. Hartvig, K. H. Andersen, J. E. Beyer, Journal of Theoretical Biology. 272, 113–122 (2011).

36. A. M. de Roos, L. Persson, Population and community ecology of ontogenetic development (Princeton University Press, 2013).

37. K. G. Srikanta Dani, U. Kodandaramaiah, Frontiers in Ecology and Evolution. 5, 1–21 (2017).

38. E. L. Charnov, S. K. M. Ernest, 167, 578–582 (2006).

39. R. W. Warne, E. L. Charnov, American Naturalist. 172 (2008), doi:10.1086/589880.

40. D. M. Anderson, J. F. Gillooly, Oikos. 130, 798–807 (2021).

41. S. M. Sogard, Bulletin of Marine Science. 60, 1129–1157 (1997).

42. J. Krause, S. P. Loader, J. McDermoi, G. D. Ruxton, Proceedings of the Royal Society B: Biological Sciences. 265, 2373–2379 (1998).

43. J. Hampton, Canadian Journal of Fisheries and Aquatic Sciences. 57, 1002–1010 (2000).

44. J. D. Arendt, Oecologia. 159, 455–461 (2009).

45. R. D. Semlitsch, Canadian Journal of Zoology. 68, 1027–1030 (1990).

46. V. H. W. Rudolf, Ecology. 89, 1650–1660 (2008).

47. T. Keren-Rotem, A. Bouskila, E. Geffen, Behavioral Ecology and Sociobiology. 59, 723–731 (2006).

48. G. W. Ferguson, S. F. Fox, Evolution. 38, 342–349 (1984).

49. J. K. Tucker, N. I. Filoramo, F. J. Janzen, The American Midland Naturalist. 141, 198–203 (1999).

50. V. H. W. Rudolf, J. Armstrong, Oecologia. 157, 675–686 (2008).

51. A. M. Boulton, G. A. Polis, The Journal of Arachnology. 27, 513–521 (1999).

52. G. Keller, G. Ribi, Oecologia. 93, 493–500 (1993).

53. S. A. H. Geritz, E. Kisdi, G. Meszena, J. A. J. Metz, Evolutionary Ecology. 12, 35–57 (1998).

54. J. Zu, M. Mimura, Y. Takeuchi, Journal of Theoretical Biology. 268, 14–29 (2011).

55. E. E. Werner, in Size Structured Populations (1988), pp. 60–81. 56. S. Diehl, P. Eklov, Ecology. 76, 1712–1726 (1995).

56. K. A. Hobson, Oecologia. 120, 134–326 (1999).

57. P. C. Chaparro-Pedraza, A. M. de Roos, Evolution. 74, 831–841 (2020).

58. P. C. Chaparro-Pedraza, A. M. de Roos, Functional Ecology. 35, 727–738 (2021).

59. P. C. Chaparro-Pedraza, A. M. de Roos, Nature Ecology and Evolution. 4, 412–418 (2020).

60. A. M. de Roos, Methods in Ecology and Evolution. 12, 383–390 (2021).

61. A. M. de Roos, in Structured-Population Models in Marine, Terrestrial, and Freshwater Systems (1997).

62. A. M. De Roos, L. Persson, Oikos. 94, 51–71 (2001).

63. R.A. Martin, D. W. Pfennig, Biological Journal of the Linnean Society. 100, 73–88 (2010).

64. H. Rundle, P. Nosil, Ecology LePers. 8, 336–352 (2005).

65. G. S. Van Doorn, P. Edelaar, F. J. Weissing, Science. 326, 1704–1707 (2009).

66. E. G. Leigh, G. J. Vermeij, Philosophical Transactions of the Royal Society B: Biological Sciences. 357, 709–718 (2002).

67. R. A. Watson et al., Evolutionary Biology. 43, 553–581 (2016).

68. S. a Levin, Contributions to Science. 7, 11–16 (2011).

69. L. Tedersoo, M. Bahram, M. Zobel, Science. 367 (2020), doi:10.1126/science.aba1223.

70. E. B. Mondor, J. A. Rosenheim, J. F. Addicoi, Functional Ecology. 22, 157–162 (2008).

71. T. E. Leonardo, E. B. Mondor, Proceedings of the Royal Society B: Biological Sciences. 273, 1079–1084 (2006).

72. G. Kunert, W. W. Weisser, Oecologia. 135, 304–312 (2003).

73. C. B. Baines, S. J. McCauley, L. Rowe, Biology LePers. 10, 10–13 (2014).

74. A. Alzate, O. Hagen, Philosophical Transactions of the Royal Society B: Biological Sciences. 379 (2024), doi:10.1098/rstb.2023.0131.

75. D. L. Rabosky et al., Nature Communications. 4, 1–8 (2013).

76. R. M. Bernstein, American Journal of Physical Anthropology. 143, 46–62 (2010).

77. J. A. Hutchings, D. W. Morris, J. A. Hutchings, D. W. Morris, Oikos. 45, 118–124 (2020).

78. N. J. Gotelli, M. Pyron, 62, 30–40 (1991).

79. A. Blanck, N. Lamouroux, Journal of Biogeography. 34, 862–875 (2007).

80. M. D. Burns, B. L. Sidlauskas, Evolution. 73, 569–587 (2019).

81. T. Takahashi, S. Koblmüller, International Journal of Evolu8onary Biology. 2011, 1–14 (2011).

82. C. J. Doenz, A.K. Krähenbühl, J. Walker, O. Seehausen, J. Brodersen, Proceedings of the Royal Society B: Biological Sciences. 286 (2019), doi:10.1098/rspb.2019.1992.

83. G. Öhlund, M. Bodin, K. A. Nilsson, S. Öhlund, K. B. Mobley, bioRxiv, 543744 (2019).

84. A. G. Hudson, P. Vonlanthen, O. Seehausen, Proceedings of the Royal Society B: Biological Sciences. 278, 58–66 (2011).

85. D. J. Fraser, L. K. Weir, L. Bernatchez, M. M. Hansen, E. B. Taylor, Heredity. 106, 404–420 (2011).

86. D. D. Heath, J. W. Heath, C. A. Bryden, R. M. Johnson, C. W. Fox, Science. 299, 1738–1740 (2003).

87. M. Ackermann, M. Doebeli, Evolution. 58, 2599–2612 (2004).

88. M. Doebeli, U. Dieckmann, 156 (2000).

89. U. Dieckmann, M. Doebeli, Nature. 400, 354–357 (1999).

90. C.H. Martin, E. J. Richards, Annual Review of Ecology, Evolution, and Systematics. 50, 569–593 (2019).

91. J. L. Payne, A. Wagner, Nature Reviews Genetics. 20, 24–38 (2019).

92. B. Maihews, R. Riera, C. C. Pedraza, C. Melian, Authorea Preprints (2024), doi:10.22541/au.173208607.77315505/v1.

93. H. Ten Brink, O. Seehausen, Proceedings of the Royal Society B: Biological Sciences. 289:202126 (2022), doi:10.1098/rspb.2021.2655.

94. H. M. Wilbur, Annual Review of Ecology and Systematics. 11, 67–93 (1980).

95. P. Chesson, 31, 343–358 (2000).

