## Supplementary material for "Life history evolution facilitates trophic diversification"

This supplementary note includes:

- Supplementary text:
    1. Ecological dynamics and derivation of the fitness gradient and curvature of the fitness landscape
    2. Analytical derivation of the ecological equilibrium
    3. Analytical derivation of conditions that enable diversification
    4. Analytical proof that the minimum productivity required for diversification occurs when the life history trait is at the optimal
    5. Model parameterization
    6. Description and parameterization of model 2 and model 3
- Figure S1

Other supplementary materials for this manuscript includes: MATLAB code of the model for numerical simulations.

#### Supplementary Information 1. Ecological dynamics and derivation of the fitness gradient and curvature of the fitness landscape

The model presented in the main text (eq. 1) can be rewritten into an equivalent model in terms of total density $N_j = A_j + J_j$, and the fraction of adults $C_j = A_j/N_j$. Changes in the variable $C_j$ therefore reflect changes in the population composition. The dynamics of the population densities, fraction of adults, and food resources thus follow

$$\frac{dN_j}{dt} = \left[\beta_j(\eta_j, \ell_j, \mathrm{F})C_j - \left(\delta_J(\ell_j)(1 - C_j) + \delta_A C_j\right)\right]N_j$$

$$\frac{dC_j}{dt} = \Phi_j(\eta_j, \ell_j, \mathrm{F})(1 - C_j) - \beta_j(\eta_j, \ell_j, \mathrm{F}){C_j}^2 - \left(\delta_A - \delta_J(\ell_j)\right)(1 - C_j)C_j,$$

$$\frac{dF_i}{dt} = \rho(F_{i\,max} - F_i) - \sum_{j=1}^{m} a_i(\eta_j)F_i\left(\gamma(1 - C_j) + C_j\right)N_j.$$

(SI1.1)

Using the adaptive dynamics framework[1,2], it is possible to derive expressions for the fitness gradient and the curvature of the fitness landscape. The relative fitness of a rare mutant population with phenotype $\mathrm{x}_j' = (\eta_j', \ell_j')$ depends on the environment set by ecomorph populations with trait values $\mathbf{x} = (\mathrm{x}_1, \dots, \mathrm{x}_m)$. Therefore, we evaluate the relative fitness of a rare mutant $\mathrm{x}_j'$ at ecological equilibrium when resident ecomorphs have traits $\mathbf{x}$:

$$W(\mathrm{x}_j', \mathbf{x}) = \beta_j\left(\eta_j', \ell_j', \mathrm{F}(\mathbf{x})\right)C_j(\ell_j') - \left(\delta_J(\ell_j')(1 - C_j(\ell_j')) + \delta_A C_j(\ell_j')\right)$$

(SI1.2)

This is because the population composition $C_j$ does depend on the life history trait but not on the feeding niche trait (see eq. SI3.16).

Under the assumption that evolution occurs via small, infrequent mutational steps, the evolutionary trajectory can be approximated by the canonical equation of adaptive dynamics[1,2]:

$$\frac{d\mathrm{x}_j}{dt} = \mathrm{M}(\mathrm{x}_j)\,\mathrm{A}(\mathrm{x}_j)\,\nabla W_j(\mathrm{x}_j, \mathbf{x})$$

(SI1.3)

In this expression, $\mathrm{M}(\mathrm{x}_j)$ is a function describing variation in the rate of the occurrence of mutations (e.g. due to variation in population size), $\mathrm{A}(\mathrm{x}_j)$ is the mutational variance-covariance matrix summarizing the distribution of mutations around the phenotype $\mathrm{x}_j$, and $\nabla W_j(\mathrm{x}_j, \mathbf{x})$ is the selection gradient, which is a two-dimensional vector whose components are:

$$\left.\frac{\partial W_j(\mathrm{x}_j',\mathbf{x})}{\partial \eta_j'}\right|_{\mathrm{x}_j'=\mathrm{x}_j} = \frac{\varepsilon C_j(\ell_j')}{\ell_j'}\sum_{i=1}^{n}\frac{\alpha\left(\theta_i-\eta_j'\right)}{\tau^2}\exp\left[\frac{-\left(\theta_i-\eta_j'\right)^2}{2\tau^2}\right]F_i(\mathbf{x}) \tag{SI1.4}$$

$$\left.\frac{\partial W_j(\mathrm{x}_j',\mathbf{x})}{\partial \ell_j'}\right|_{\mathrm{x}_j'=\mathrm{x}_j} = \frac{\varepsilon}{\ell_j'}\left(\frac{\partial C_j}{\partial \ell_j'}-\frac{C_j(\ell_j')}{\ell_j'}\right)\sum_{i=1}^{n}a_i(\eta_j')F_i(\mathbf{x})-\frac{\nu}{\ell_j'}\left(\frac{\partial C_j}{\partial \ell_j'}-\frac{C_j(\ell_j')}{\ell_j'}\right) + \delta_{\max}e^{-\ell_j'}\left(1-C_j(\ell_j')+\frac{\partial C_j}{\partial \ell_j'}\right)-\delta_A\frac{\partial C_j}{\partial \ell_j'} \tag{SI1.5}$$

Diversification occurs through a process of evolutionary branching when directional selection in the feeding niche trait halts, if the niche trait value at this point corresponds to a minimum of the fitness landscape [3,4]. The curvature of the fitness function therefore determines whether an ecomorph splits into two different ecomorphs. Hence, the second derivative of the first component of the fitness function with respect to $\eta_j'$ evaluated at the ecological equilibrium when ecomorphs have traits $\mathbf{x}$ allows us to determine whether a diversification event occurs

$$\left.\frac{\partial^2 W_j(\mathrm{x}_j',\mathbf{x})}{\partial {\eta_j'}^2}\right|_{\mathrm{x}_j'=\mathrm{x}_j} = \frac{\varepsilon C_j(\ell_j')}{\ell_j'}\sum_{i=1}^{n}\frac{\alpha\ \left(\left(\theta_i-\eta_j'\right)^2-\tau^2\right)}{\tau^4}\exp\left[\frac{-\left(\theta_i-\eta_j'\right)^2}{2\tau^2}\right]F_i(\mathbf{x}). \tag{SI1.6}$$

**References**


1. Dieckmann, U. & Law, R. *J. Math. Biol.* **34**, 579–612 (1996).
2. Leimar, O. *Evol. Ecol. Res.* **11**, 191–208 (2009).
3. Dieckmann, U. (Cambridge University Press., 2004). doi:10.1017/CBO9781139342179
4. Geritz, S. A. H. & Kisdi, E. *Proc. R. Soc. B Biol. Sci.* **267**, 1671–1678 (2000).

### Supplementary Information 2. Analytical derivation of the ecological equilibrium

Here, we derive expressions for the ecological equilibrium when one population with total density $N$, fraction of adults $C$, feeding niche trait $\eta$, and offspring size $\ell$ colonizes an environment with two different food resources, $F_1$ and $F_2$. The analytical investigation is restricted to the simplest case, in which $F_{1max} = F_{2max} = F_{max}$.

From eq. SI1.1 in SI1, the population density, fraction of juveniles and resources satisfy the following equations at ecological equilibrium

$$0 = C\left(\varepsilon(a_1(\eta)F_1 + a_2(\eta)F_2) - v\right)/\ell - \delta_J(\ell)(1-C) - \delta_A C) \quad \text{(SI2.1)}$$

$$0 = \Phi(\eta, \ell, F_1, F_2)(1-C) - \left(\varepsilon(a_1(\eta)F_1 + a_2(\eta)F_2) - v\right)C^2/\ell - (\delta_A - \delta_J(\ell))(1-C)C \quad \text{(SI2.2)}$$

$$0 = \rho(F_{max} - F_1) - a_1(\eta)F_1\left(\gamma(1-C) + C\right)N \quad \text{(SI2.3)}$$

$$0 = \rho(F_{max} - F_2) - a_2(\eta)F_2\left(\gamma(1-C) + C\right)N \quad \text{(SI2.4)}$$

Equations SI2.1, SI2.3, SI2.4 can be reformulated to obtain

$$0 = E(a_1(\eta)F_1 + a_2(\eta)F_2) - G \quad \text{(SI2.5)}$$

$$0 = \rho(F_{max} - F_1) - a_1(\eta)F_1 IN \quad \text{(SI2.6)}$$

$$0 = \rho(F_{max} - F_2) - a_2(\eta)F_2 IN \quad \text{(SI2.7)}$$

where $E = \varepsilon C/\ell$, $G = vC/\ell + \delta_A C + \delta_J(1-C)$, and $I = \gamma(1-C) + C$.

Equations (SI2.6) and (SI2.7) give

$$F_1 = \frac{\rho F_{max}}{\rho + a_1(\eta)IN} \text{ and } F_2 = \frac{\rho F_{max}}{\rho + a_2(\eta)IN} \quad \text{(SI2.8)}$$

By substituting $F_1$ and $F_2$ in equation (SI2.5), an equation for $N$ is obtained:

$$0 = E\left(a_1(\eta)\frac{\rho F_{max}}{\rho + a_1(\eta)IN} + a_2(\eta)\frac{\rho F_{max}}{\rho + a_2(\eta)IN}\right) - G$$

which is equivalent to the quadratic equation

$$0 = \alpha_N N^2 + \beta_N N + \gamma_N \quad \text{(SI2.9)}$$

where

$$\alpha_N = GI^2 a_1(\eta)a_2(\eta) \quad \text{(SI2.10)}$$

$$\beta_N = \rho GI(a_1(\eta) + a_2(\eta)) - 2EI\rho F_{max}a_1(\eta)a_2(\eta) \quad \text{(SI2.11)}$$

$$\gamma_N = \rho^2 G - E\rho^2 F_{max}(a_1(\eta) + a_2(\eta)) \quad \text{(SI2.12)}$$

The two solutions of equation (SI2.9) are

$$N^*(\eta) = \frac{-\beta_N \pm \sqrt{\Delta_N}}{2\alpha_N}$$

where $\Delta_N = \beta_N^2 - 4\alpha_N\gamma_N = G^2\rho^2 I^2(a_1(\eta) - a_2(\eta))^2 + 4E^2I^2\rho^2F_{max}^2 a_1(\eta)^2 a_2(\eta)^2$. Only when $\sqrt{\Delta_N}$ is added to $-\beta_N$ the population density is positive, therefore, the only biologically relevant solution is

$$N^*(\eta) = \frac{-\beta_N + \sqrt{\Delta_N}}{2\alpha_N} \tag{SI2.13}$$

By substituting $N^*$ in equations (SI2.8), it can be obtained

$$F_1^*(\eta) = \frac{2a_2(\eta)\rho F_{max}GI}{\sqrt{\rho^2(4B^2 + G^2I^2(a_2(\eta) - a_1(\eta))^2)} + \rho(2B + GI(a_2(\eta) - a_1(\eta)))} \tag{SI2.14}$$

$$F_2^*(\eta) = \frac{2a_1(\eta)\rho F_{max}GI}{\sqrt{\rho^2(4B^2 + G^2I^2(a_2(\eta) - a_1(\eta))^2)} + \rho(2B - GI(a_2(\eta) - a_1(\eta)))}, \tag{SI2.15}$$

where $B = EIF_{max}a_1(\eta)a_2(\eta)$.

By replacing the maturation function $\Phi(\eta, \ell, F_1, F_2)$ into equation (SI2.3) one obtains:

$$0 = \left( \frac{\varepsilon\gamma(a_1(\eta)F_1 + a_2(\eta)F_2) - v - \delta_J(\ell)}{1 - \left(\frac{\ell}{w}\right)^{1 - \frac{\delta_J(\ell)}{\varepsilon\gamma(a_1(\eta)F_1 + a_2(\eta)F_2) - v}}} \right)(1-C) - (\varepsilon(a_1(\eta)F_1 + a_2(\eta)F_2) - v)\frac{C^2}{\ell} - (\delta_J - \delta_A)(1-C)C$$

From equation (SI2.5) we have that in the ecological equilibrium $a_1(\eta)F_1 + a_2(\eta)F_2 = \frac{\ell G}{\varepsilon C}$. Therefore, this equation can be reformulated as

$$0 = \left( \frac{\gamma\frac{G\ell}{C} - v - \delta_J(\ell)}{1 - \left(\frac{\ell}{w}\right)^{1 - \frac{\delta_J(\ell)}{\gamma\frac{\ell G}{C} - v}}} \right)(1 - C) - \left(\frac{G\ell}{C} - v\right)\frac{C^2}{\ell} - (\delta_A - \delta_J(\ell))(1 - C)C \tag{SI2.16}$$

Unfortunately, this expression cannot be solved analytically due to the complexity of the maturation function. Therefore, we cannot obtain an analytical expression for the fraction of adults $C^*$ in the ecological equilibrium. However, from equation SI2.16 it is possible to infer that the fraction of adults $C^*$ in the ecological equilibrium does not depend on the trait $\eta$.

### Supplementary Information 3. Analytical derivation of conditions that enable diversification

Here, I derive the conditions that enable diversification when one population with total density $N$, fraction of adults $C$, feeding niche trait $\eta$, and offspring size $\ell$ colonizes an environment with two different food resources, $F_1$ and $F_2$. Because diversification can only occur in the feeding niche trait $\eta$, we study the evolution of this trait while offspring size $\ell$ is kept fixed.

For diversification to occur, a local minimum of the fitness landscape must be an attractor of the evolutionary dynamics. Thus, two conditions must be fulfilled for diversification to occur: 1) the mean phenotype of the population at evolutionary equilibrium corresponds to a local fitness minimum such that selection is disruptive, and 2) this equilibrium is an attractor of the evolutionary dynamics.

To investigate these conditions, we use the adaptive dynamics framework, which assumes that evolution occurs much more slowly than the ecological dynamics. Under this assumption, the system reaches its ecological attractor in between two mutations. Using the analytical expressions for the ecological equilibrium obtained in Supplementary information 3, we will investigate the fitness landscape of a rare mutant in the environment set by the population with a trait value $\eta$. The optimal niche trait values to feed on resource 1 and resource 2 are $\theta_1$ and $\theta_2$, respectively, and their distance in the trait axis equals $D$. Without a loss of generality, it can be assumed that $\theta_1 = 0$ and $\theta_2 = D$.

**1. When does a population experience disruptive selection?** One of the necessary conditions for diversification is that a population experiences disruptive selection. This occurs when the population is in an evolutionary equilibrium that corresponds to a fitness minimum, hence the curvature of the fitness landscape around its trait value is positive. In this section, I investigate when this condition is satisfied by analysing the fitness landscape of a rare mutant with trait value $\eta'$, in the environment set by a population with a trait value $\eta^* = \frac{D}{2}$.

First, we show that $\eta^* = \frac{D}{2}$ is an evolutionary equilibrium. To do so, we calculate the selection gradient using eq. (SI1.2):

$$\begin{aligned}
\frac{\partial W}{\partial \eta'}(\eta',\eta) &= E\left(\frac{\partial a_1}{\partial \eta'}F_1(\eta) + \frac{\partial a_2}{\partial \eta'}F_2(\eta)\right) \\
&= E\left(-\frac{\eta}{\tau^2}(a_1(\eta)F_1(\eta) + a_2(\eta)F_2(\eta)) + \frac{D}{\tau^2}a_2(\eta)F_2(\eta)\right) \\
&= E\left(-\frac{\eta}{\tau^2}\left(\frac{G}{E}\right) + \frac{D}{\tau^2}a_2(\eta)F_2(\eta)\right) \\
&= -\frac{\eta G}{\tau^2} + E\frac{D}{\tau^2}a_2(\eta)F_2(\eta). \qquad \text{(SI3.1)}
\end{aligned}$$

Between step two and three, we use $a_1(\eta)F_1 + a_2(\eta)F_2 = \frac{G}{E}$ from eq. (SI2.5).

When $\eta^* = \frac{D}{2}$, $F_1^*\left(\frac{D}{2}\right) = F_2^*\left(\frac{D}{2}\right)$, hence, using equation (SI2.5), one gets

$$a_2\left(\frac{D}{2}\right)F_2\left(\frac{D}{2}\right) = \frac{G}{2E}. \qquad \text{(SI3.2)}$$

Substituting equation (SI3.2) in equation (SI3.1), it is posible to show that at $\eta^* = \frac{D}{2}$ the selection gradient is always zero, and hence that this trait is an evolutionary equilibrium for any $D > 0$:

$$\frac{\partial W}{\partial \eta'}\left(\frac{D}{2}, \frac{D}{2}\right) = -\frac{DG}{2\tau^2} + E\frac{DG}{2\tau^2 E} = 0. \quad \text{(SI3.3)}$$

Then, the curvature of the fitness landscape is calculated:

$$\begin{aligned}\frac{\partial^2 W}{\partial \eta'^2}(\eta', \eta) &= E\left(\frac{\partial^2 a_1}{\partial \eta'^2}F_1(\eta) + \frac{\partial^2 a_2}{\partial \eta'^2}F_2(\eta)\right) \\ &= E\left(\frac{1}{\tau^2}a_1(\eta)\left(\frac{\eta^2}{\tau^2} - 1\right)F_1(\eta) + \frac{1}{\tau^2}a_2(\eta)\left(\frac{(\eta - D)^2}{\tau^2} - 1\right)F_2(\eta)\right) \\ &= \frac{1}{\tau^2}\left(\left(\frac{\eta^2}{\tau^2} - 1\right)G + E\frac{D}{\tau^2}a_2(\eta)F_2(\eta)(D - 2\eta)\right). \quad \text{(SI3.4)}\end{aligned}$$

Finally, we evaluate the curvature of the fitness landscape at $\eta^* = \frac{D}{2}$

$$\frac{\partial^2 W}{\partial \eta'^2}\left(\frac{D}{2}, \frac{D}{2}\right) = \frac{G}{\tau^2}\left(\frac{D^2}{4\tau^2} - 1\right) \quad \text{(SI3.5)}$$

The curvature of the fitness landscape for $\eta^* = \frac{D}{2}$ is positive, i.e. $\frac{\partial^2 W}{\partial \eta'^2}(\frac{D}{2}, \frac{D}{2}) > 0$, and thus selection is disruptive in the evolutionary equilibrium $\eta^* = \frac{D}{2}$ when

$$D > 2\tau \quad \text{(SI3.6)}$$

**2. When does a population evolve toward the evolutionary equilibrium where selection is disruptive?** The second necessary condition for diversification is that a population evolves toward the evolutionary equilibrium where selection is disruptive. This occurs when this equilibrium is an evolutionary attractor. To determine when the equilibrium $\eta^* = \frac{D}{2}$ is an attractor, we investigate its stability.

An evolutionary equilibrium $\eta^*$ is stable if $\frac{\partial}{\partial \eta}\left(\frac{\partial W}{\partial \eta'}(\eta, \eta)\right) < 0$. Hence, we first differentiate equation (SI3.1) with respect to $\eta$.

$$\begin{aligned}\frac{\partial}{\partial \eta}\left(\frac{\partial W}{\partial \eta'}(\eta', \eta)\right) &= -\frac{G}{\tau^2} + E\frac{D}{\tau^2}\left(\frac{\partial a_2}{\partial \eta}(\eta)F_2(\eta) + a_2(\eta)\frac{\partial F_2}{\partial \eta}(\eta)\right) \\ &= -\frac{G}{\tau^2} + E\frac{D}{\tau^2}\left(\frac{D - \eta}{\tau^2}a_2(\eta)F_2(\eta) + a_2(\eta)\frac{\partial F_2}{\partial \eta}(\eta)\right)\end{aligned} \quad \text{(SI3.7)}$$

Next, we calculate $\frac{\partial F_2}{\partial \eta}(\eta)$ by derivating eq. (SI2.8) with respect to $\eta$

$$\begin{aligned}\frac{\partial F_2}{\partial \eta} &= \frac{-\rho F_{max}\left(\frac{\partial a_2}{\partial \eta}IN + a_2 I\frac{\partial N}{\partial \eta}\right)}{(\rho + a_2 IN)^2} \\ &= -F_2(\eta)\frac{\frac{\partial a_2}{\partial \eta}IN + a_2 I\frac{\partial N}{\partial \eta}}{\rho + a_2 IN}\end{aligned}$$

$\frac{\partial N}{\partial \eta}$ can be obtained by derivating eq. (SI2.10), (SI2.11), (SI2.12), and (SI2.13):

$$\frac{\partial N^*}{\partial \eta} = \frac{2\alpha_N\left(-\frac{\partial \beta_N}{\partial \eta} + \frac{\partial}{\partial \eta}(\sqrt{\Delta_N})\right) - \left(-\beta_N + \sqrt{\Delta_N}\right)\frac{\partial \alpha_N}{\partial \eta}}{4\alpha_N^2} \tag{SI3.8}$$

where

$$\frac{\partial \alpha_N}{\partial \eta} = \frac{GI^2}{\tau^2}a_1(\eta)a_2(\eta)(D-2\eta) \tag{SI3.9}$$

$$\frac{\partial \beta_N}{\partial \eta} = -\frac{\eta}{\tau^2}\rho GIa_1(\eta) - \frac{\eta - D}{\tau^2}\rho GIa_2(\eta) - \frac{2(D-2\eta)}{\tau^2}a_1(\eta)a_2(\eta)EI\rho F_{max} \tag{SI3.10}$$

$$\begin{aligned}\frac{\partial \sqrt{\Delta_N}}{\partial \eta} =& \frac{1}{\sqrt{\Delta_N}}\left[G^2\rho^2I^2(a_1(\eta) - a_2(\eta))\left(\frac{-\eta}{\tau^2}a_1(\eta) + \frac{\eta - D}{\tau^2}a_2(\eta)\right)\right. \\ &\left. +4E^2I^2\rho^2F_{max}^2a_1(\eta)^2a_2(\eta)^2\left(\frac{D-2\eta}{\tau^2}\right)\right]\end{aligned} \tag{SI3.11}$$

(SI3.12)

Subsequently, we evaluate $\frac{\partial}{\partial \eta}\left(\frac{\partial W}{\partial \eta'}(\eta', \eta)\right)$ at $\eta^* = \frac{D}{2}$

$$\frac{\partial}{\partial \eta}\left(\frac{\partial W}{\partial \eta'}(\eta', \eta)\right)(\frac{D}{2}) = -\frac{G}{\tau^2} + \frac{GD^2}{4\tau^4} + E\frac{D}{\tau^2}a_2(\frac{D}{2})\frac{\partial F_2(\frac{D}{2})}{\partial \eta}. \tag{SI3.13}$$

Evaluating $\frac{\partial N^*}{\partial \eta}(\eta)$ at $\eta^* = \frac{D}{2}$, one gets

$$\frac{\partial N^*}{\partial \eta}(\frac{D}{2}) = 0 \tag{SI3.14}$$

because $\frac{\partial \alpha_N}{\partial \eta}(\frac{D}{2}) = 0$, $\frac{\partial \beta_N}{\partial \eta}(\frac{D}{2}) = 0$, and $\frac{\partial \sqrt{\Delta}}{\partial \eta}(\frac{D}{2}) = 0$.

Evaluating $\frac{\partial F_2}{\partial \eta}(\eta)$ at $\eta^* = \frac{D}{2}$, one gets

$$\begin{aligned}\frac{\partial F_2}{\partial \eta}(\frac{D}{2}) &= -F_2\frac{D}{2\tau^2}\frac{a_2(\eta)IN^*}{\rho + a_2(\eta)IN^*} \\ &= -\frac{G}{2Ea_2(\eta)}\frac{D}{2\tau^2}\frac{a_2(\eta)IN^*}{\rho + a_2(\eta)IN^*} \\ &= -\frac{GID}{4\tau^2E}\frac{N^*}{\rho + a_2(\eta)IN^*} \\ &= -\frac{GID}{4\tau^2E}\left(\frac{1}{Ia_2(\eta)} - \frac{G}{2EIF_{max}a_2(\eta)^2}\right) \\ &= -\frac{GD}{4\tau^2E}\left(\frac{1}{a_2(\eta)} - \frac{G}{2EF_{max}a_2(\eta)^2}\right)\end{aligned} \tag{SI3.15}$$

because

$$\begin{aligned}N^*(\frac{D}{2}) &= \frac{-2\rho GIa_2(\frac{D}{2}) + 2EI\rho F_{max}a_2(\frac{D}{2})^2 + 2EI\rho F_{max}a_2(\frac{D}{2})^2}{2GI^2a_2(\frac{D}{2})^2} \\ &= \frac{-\rho G + 2E\rho F_{max}a_2(\frac{D}{2})}{GIa_2(\frac{D}{2})} \\ &= \frac{-\rho}{Ia_2(\frac{D}{2})} + \frac{2E\rho F_{max}}{GI}\end{aligned} \tag{SI3.16}$$

Replacing eq. (SI3.15) in eq. (SI3.13), it yields

$$
\begin{aligned}
\frac{\partial}{\partial \eta}\left(\frac{\partial W}{\partial \eta'}(\eta,\eta)\right)(\frac{D}{2}) &= -\frac{G}{\tau^2} + \frac{GD^2}{4\tau^4} \\
&+ E\frac{D}{\tau^2}a_2(\frac{D}{2})\left(-\frac{GD}{4\tau^2 E}\left(\frac{1}{a_2(\frac{D}{2})} - \frac{G}{2EF_{max}a_2(\frac{D}{2})^2}\right)\right) \\
&= -\frac{G}{\tau^2} + \frac{GD^2}{4\tau^4} + \frac{GD^2}{4\tau^4}\left(-1 + \frac{G}{2EF_{max}a_2(\frac{D}{2})}\right) \\
&= -\frac{G}{\tau^2} + \frac{G^2D^2}{8\tau^4 EF_{max}a_2(\frac{D}{2})} \\
&= -\frac{G}{\tau^2} + \frac{G^2D^2}{8\tau^4 EF_{max}\alpha}e^{\frac{D^2}{8\tau^2}} \qquad \text{(SI3.17)}
\end{aligned}
$$

For $\eta^* = \frac{D}{2}$ to be an evolutionary attractor, and thus for selection to drive the feeding niche trait towards this evoutionary equilibrium, $\frac{\partial}{\partial \eta}\left(\frac{\partial W}{\partial \eta'}(\eta,\eta)\right)(\frac{D}{2}) < 0$. This is possible when:

$$
\begin{aligned}
-\frac{G}{\tau^2} + \frac{G^2D^2}{8\tau^4 EF_{max}\alpha}e^{\frac{D^2}{8\tau^2}} &< 0 \\
\frac{GD^2}{8\tau^2 EF_{max}\alpha}e^{\frac{D^2}{8\tau^2}} &< 1 \qquad \text{(SI3.18)}
\end{aligned}
$$

In conclusion, diversification occurs at the evolutionary equilibrium $\eta^* = \frac{D}{2}$ when the two following conditions are met:

$$
\begin{aligned}
2\tau &< D \qquad \text{(SI3.19)} \\
\frac{GD^2}{8\tau^2 E\alpha}e^{\frac{D^2}{8\tau^2}} &< F_{max}. \qquad \text{(SI3.20)}
\end{aligned}
$$

The second condition can be rewritten in terms of the productivity of the system, i.e. $P = \rho(F_{1max} + F_{2max}) = 2\rho F_{max}$:

$$
\frac{\rho GD^2}{4\tau^2 E\alpha}e^{\frac{D^2}{8\tau^2}} < P. \qquad \text{(SI3.21)}
$$

### Supplementary Information 4. Analytical proof that the minimum productivity required for diversification occurs when the life history trait is at the optimal

Here, I show that the threshold of productivity required for diversification takes the lowest value when the selection gradient of the life history trait (offspring size) vanishes, or in other words, when this trait is equal to the optimal. As in the previous sections, we consider a population with total density $N$, fraction of adults $C$, and phenotype $x = (\eta, \ell)$, where $\eta$ is the feeding niche trait, and $\ell$ is the offspring size. The population colonizes an environment with two different food resources, $F_1$ and $F_2$. Using the adaptive dynamics framework, which assumes that evolution occurs much more slowly than the ecological dynamics, and thus that the system reaches its ecological attractor in between two mutations, I derive an expression for the offspring size when the selection gradient vanishes. Then, I calculate the minimum productivity required for diversification as a function of the offspring size.

**1. Optimal offspring size** From eq. SI1.5, the selection gradient of a rare mutant with offspring size $\ell'$ is

$$\frac{\partial W}{\partial \ell'}(x', x) = \frac{\varepsilon}{\ell'}\left(\frac{\partial C}{\partial \ell'} - \frac{C}{\ell'}\right)(a_1(\eta)F_1 + a_2(\eta)F_2) - \frac{v}{\ell'}\left(\frac{\partial C}{\partial \ell'} - \frac{C}{\ell'}\right) + \delta_{max}e^{-\ell'}\left(1 - C + \frac{\partial C}{\partial \ell'}\right) - \delta_A\frac{\partial C}{\partial \ell'} \tag{SI4.1}$$

Because in the ecological equilibrium $a_1(\eta)F_1 + a_2(\eta)F_2 = \frac{G}{E}$ (see eq. SI2.5), we have

$$\begin{aligned}\frac{\partial W}{\partial \ell'}(x', x) &= \frac{G}{C}\frac{\partial C}{\partial \ell'} - \frac{G}{\ell'} - \frac{V}{\ell}\frac{\partial C}{\partial \ell'} + \frac{VC}{\ell'^2} + \delta_{max}e^{-\ell'}(1 - C) + \delta_{max}e^{-\ell'}\frac{\partial C}{\partial \ell'} - \delta_A\frac{\partial C}{\partial \ell'} \\ &= \left(\frac{G}{C} - \frac{V}{\ell'} + \delta_{max}e^{-\ell'} - \delta_A\right)\frac{\partial C}{\partial \ell'} - \frac{G}{\ell'} + \frac{VC}{\ell'^2} + \delta_{max}e^{-\ell'}(1 - C). \end{aligned} \tag{SI4.2}$$

Replacing $G = \frac{VC}{\ell} + \delta_A C + \delta_{max}e^{-\ell}(1 - C)$, we get

$$\begin{aligned}\frac{\partial W}{\partial \ell'}(x', x) =& \left(\frac{1}{C}\left(\frac{VC}{\ell'} + \delta_A C + \delta_{max}e^{-\ell'}(1 - C)\right) - \frac{V}{\ell'} + \delta_{max}e^{-\ell'} - \delta_A\right)\frac{\partial C}{\partial \ell'} \\ & - \frac{1}{\ell'}\left(\frac{VC}{\ell'} + \delta_A C + \delta_{max}e^{-\ell'}(1 - C)\right) + \frac{VC}{\ell'^2} + \delta_{max}e^{-\ell'}(1 - C) \\ =& \frac{\delta_{max}e^{-\ell'}}{C}\frac{\partial C}{\partial \ell'} + \frac{1}{\ell'}\left(-\delta_A C + \delta_{max}e^{-\ell'}(1 - C)(\ell' - 1)\right)\end{aligned} \tag{SI4.3}$$

The trait value at which the selection gradient equals zero is the optimal value for the offspring size. Therefore, this trait value can be found by solving $\ell$ from this equation:

$$0 = \frac{\delta_{max}e^{-\ell}}{C}\frac{\partial C}{\partial \ell} + \frac{1}{\ell}\left(-\delta_A C + \delta_{max}e^{-\ell}(1 - C)(\ell - 1)\right). \tag{SI4.4}$$

However, it is not possible to obtain an analytical solution for this equation because we do not have an analytical expression for $C$ and $\frac{\partial C}{\partial \ell}$ (see SI2.16).

**2. Minimum productivity for diversification** From eq. SI3.21, we have that the threshold of productivity required for diversification $P_{min}$ is

$$P_{min} = \frac{\rho G D^2}{4\tau^2 E\alpha} e^{\frac{D^2}{8\tau^2}} \tag{SI4.5}$$

Replacing the terms $G$ and $E$ (see eq. SI2.5), $P_{min}$ can be expressed as

$$\begin{aligned} P_{min} &= \frac{\ell}{\varepsilon C}\left(\frac{vC}{\ell} + \delta_A C + \delta_{max} e^{-\ell}(1-C)\right)\frac{\rho D^2}{4\tau^2\alpha} e^{\frac{D^2}{8\tau^2}} \\ &= \left(\frac{v}{\varepsilon} + \frac{\ell\delta_A}{\varepsilon} + \frac{\ell\delta_{max}e^{-\ell}}{\varepsilon C} - \frac{\ell\delta_{max}e^{-\ell}}{\varepsilon}\right)\frac{\rho D^2}{4\tau^2\alpha} e^{\frac{D^2}{8\tau^2}} \end{aligned} \tag{SI4.6}$$

To understand how $P_{min}$ depends on $\ell$, we derivate this expression with respect to $\ell$

$$\frac{\partial P_{min}}{\partial \ell} = \left(\frac{\delta_A}{\varepsilon} - \frac{\delta_{max}e^{-\ell}}{\varepsilon} + \frac{\ell\delta_{max}e^{-\ell}}{\varepsilon} + \frac{\delta_{max}e^{-\ell}}{\varepsilon C} - \frac{\ell\delta_{max}e^{-\ell}}{\varepsilon C} - \frac{\ell\delta_{max}e^{-\ell}}{\varepsilon C^2}\frac{\partial C}{\partial \ell}\right)\frac{\rho D^2}{4\tau^2\alpha} e^{\frac{D^2}{8\tau^2}} \tag{SI4.7}$$

Then, we set this function to zero to find the minimum

$$\begin{aligned} 0 &= \delta_A C - \delta_{max}e^{-\ell}C + \ell\delta_{max}e^{-\ell}C + \delta_{max}e^{-\ell} - \ell\delta_{max}e^{-\ell} - \frac{\ell\delta_{max}e^{-\ell}}{C}\frac{\partial C}{\partial \ell} \\ &= \delta_A C - \delta_{max}e^{-\ell}(1-C)(\ell-1) - \frac{\ell\delta_{max}e^{-\ell}}{C}\frac{\partial C}{\partial \ell} \\ &= \frac{1}{\ell}\left(-\delta_A C + \delta_{max}e^{-\ell}(1-C)(\ell-1)\right) + \frac{\delta_{max}e^{-\ell}}{C}\frac{\partial C}{\partial \ell} \end{aligned} \tag{SI4.8}$$

Therefore, the threshold of minimum productivity required for diversification takes its lowest value when the offspring size takes the value of $\ell$ that satisfies this equation. Because this expression is equivalent to the expression for the optimal offspring size in equation (SI4.4), we can affirm that the threshold of productivity required for diversification takes its lowest value when the offspring size is at the optimal value.

### Supplementary Information 5. Model parameterization

The model presented in the main text is based on previous models of ecological diversification (e.g. ref [1–3]). In particular, we extend the stage-structured model studied by Chaparro-Pedraza[3] and adopt the values for the environmental parameters and demographic parameters independent of stage structure (i.e. $\alpha$, $\tau$, $\varepsilon$, $\nu$) used in that study. Some of these values are key for diversification and thus selected accordingly, for example, $\tau$ and the distance between the two optima to feed on the food resources $D$ must satisfy the condition $D > 2\tau$ to ensure that selection is disruptive at the trait value in between the two optima (see SI3, eq. SI3.3). Additionally, for the stage-dependent demographic parameters we adopt values that broadly represent organisms in nature. The general rule across animal taxa is that larger organisms have higher survival[4–15] and higher foraging capacity[16–18] than smaller conspecifics. Hence, stage-specific parameters are assigned values that results in a higher survival and foraging capacity of adults than juveniles (i.e. $\delta_A < \delta_J$ and $\gamma < 1$). $\gamma = 0.5$ is adopted as the default value for the ingestion rate of juveniles, similar to values previously used in similar stage-structured models (e.g. ref [19]). Parameter values are summarized in table S1.

Table SI5.1. Variables and parameters

| Variable or parameter | Symbol | Value | Units |
|---|---|---|---|
| Variables | | | |
| Feeding niche trait | $\eta$ | Evolving trait | - |
| Offspring size | $\ell$ | Evolving trait | g |
| Individual density | $N_j$ | - | $L^{-1}$ |
| Adult fraction | $C_j$ | - | - |
| Food resource density | $F_i$ | - | g $L^{-1}$ |
| Environmental parameters | | | |
| Renewal rate of resources | $\rho$ | 0.01 | (unit of time)$^{-1}$ |
| Total productivity of the habitat | $P$ | 0.4 (by default) varied in fig. 4 | g $L^{-1}$ (unit of time)$^{-1}$ |
| Number of resources | $n$ | 10 | - |
| Distance between optima to feed on the resources | $D$ | 1 | - |
| Demographic parameters | | | |
| Maximum attack rate | $\alpha$ | 0.2 | L (unit of time)$^{-1}$ |
| Width of the Gaussian curve describing the degree of specialization to successfully attack a food resource | $\tau$ | 1/3 | - |
| Assimilation efficiency | $\varepsilon$ | 0.6 | - |
| Metabolic cost | $\nu$ | 0.01 | (weight)*(unit of time)$^{-1}$ |
| Juvenile foraging capacity factor | $\gamma$ | 0.5 | - |
| Adult mortality rate | $\delta_A$ | 0.01 | (unit of time)$^{-1}$ |
| Maximum juvenile mortality rate | $\delta_{\max}$ | 0.01 5 (by default) varied in fig. 4 | (unit of time)$^{-1}$ |
| Size at maturation | $w$ | 10 | g |
| Evolutionary parameters | | | |
| Mutation rate of the feeding niche trait | $\mu_\eta$ | $10^{-3}$ | - |
| Mutation rate of the offspring size | $\mu_\ell$ | $10^{-3}$ (by default) varied in fig. 3 | - |
| Variance of the trait offspring distribution (feeding niche trait) | $\sigma_\eta^2$ | 0.01 | - |
| Variance of the trait offspring distribution (offspring size) | $\sigma_\ell^2$ | 0.01 | - |

#### References


1. Chaparro-Pedraza, C., Roth, G. & Melian, C. Ecological diversification in sexual and asexual lineages. *bioRxiv* 2024.03.06.583698 (2024) doi:10.1101/2024.03.06.583698.
2. Chaparro-Pedraza, P. C., Roth, G. & Seehausen, O. The enrichment paradox in adaptive radiations: Emergence of predators hinders diversification in resource rich environments. *Ecol. Lett.* **25**, 802–813 (2022).
3. Chaparro-Pedraza, P. C. Differential stage-specific mortality as a mechanism for diversification. *Am. Nat.* **(in press)**, (2024).
4. Sogard, S. M. Size selective mortality in the juvenile stages of teleost fishes: a review. *Bull. Mar. Sci.* **60**, 1129–1157 (1997).
5. Krause, J., Loader, S. P., McDermott, J. & Ruxton, G. D. Refuge use by fish as a function of body length-related metabolic expenditure and predation risks. *Proc. R. Soc. B Biol. Sci.* **265**, 2373–2379 (1998).
6. Boulton, A. M. & Polis, G. A. Phenology and Life History of the Desert Spider, Diguetia mojavea (Araneae , Diguetidae). *J. Arachnol.* **27**, 513–521 (1999).
7. Keller, G. & Ribi, G. Fish predation and offspring survival in the prosobranch snail Viviparus ater. *Oecologia* **93**, 493–500 (1993).
8. Hampton, J. Natural mortality rates in tropical tunas: size really does matter. *Can. J. Fish. Aquat. Sci.* **57**, 1002–1010 (2000).
9. Arendt, J. D. Influence of sprint speed and body size on predator avoidance in New Mexican spadefoot toads (Spea multiplicata). *Oecologia* **159**, 455–461 (2009).
10. Semlitsch, R. D. Effects of body size, sibship, and tail injury on the susceptibility of tadpoles to dragonfly predation. *Can. J. Zool.* **68**, 1027–1030 (1990).
11. Rudolf, V. H. W. Impact of Cannibalism on Predator-Prey Dynamics : Size-Structured Interactions and Apparent Mutualism. *Ecology* **89**, 1650–1660 (2008).
12. Keren-Rotem, T., Bouskila, A. & Geffen, E. Ontogenetic habitat shift and risk of cannibalism in the common chameleon (Chamaeleo chamaeleon). *Behav. Ecol. Sociobiol.* **59**, 723–731 (2006).
13. Ferguson, G. W. & Fox, S. F. Annual Variation of Survival Advantage of Large Juvenile Side-Blotched Lizards, Uta stansburiana: Its Causes and Evolutionary Significance. *Evolution (N. Y).* **38**, 342–349 (1984).
14. Tucker, J. K., Filoramo, N. I. & Janzen, F. J. Size-biased mortality due to predation in a nesting freshwater turtle, Trachemys scripta. *Am. Midl. Nat.* **141**, 198–203 (1999).
15. Rudolf, V. H. W. & Armstrong, J. Emergent impacts of cannibalism and size refuges in prey on intraguild predation systems. *Oecologia* **157**, 675–686 (2008).
16. Gribbin, S. D. & Thompson, D. J. Asymmetric intraspecific competition among larvae of the damselfly Ischnura elegans (Zygoptera: Coenagrionidae). *Ecol. Entomol.* **15**, 37–42 (1990).
17. McPeek, M. A. & Crowley, P. H. The effects of density and relative size on the aggressive behaviour, movement and feeding of damselfly larvae (Odonata: Coenagrionidae). *Anim. Behav.* **35**, 1051–1061 (1987).
18. Nakayama, S. & Fuiman, L. A. Body size and vigilance mediate asymmetric interference competition for food in fish larvae. *Behav. Ecol.* **21**, 708–713 (2010).
19. de Roos, M. Dynamic population stage structure due to juvenile – adult asymmetry stabilizes complex ecological communities. *Proc. Natl. Acad. Sci.* **118**, 1–8 (2021).

#### Supplementary Information 6. Description and parameterization of model 2 and 3

In this supplementary note, we describe the size-structured population models used to test the generality of our findings. The core part of these models is the description of the individual behavior of the consumer ecomorphs, that is, individual feeding, growth, development, reproduction and mortality, as a function of the current state of both the environment (e.g. resource availability) and the individual itself (i.e. its body size).

##### *Model 2: Diversification and evolution of maturation size*

The model follows the food web bioenergetic approach for size-structured populations introduced by Hartvig[1]. Here we only provide a concise synopsis of the model. We consider an ecomorph population $j$ to be composed of individuals characterized by their feeding niche trait $\eta_j$, body size $s$ and body size at maturation $s_m$. The rate at which an individual encounters food depends on $\eta_j$, $s$ and the densities of the resources $\mathrm{F} = (F_1 \dots F_n)$, such that

$$c_{j,s}(\eta_j, s, \mathrm{F}) = \sum_{i=1}^{n} A_{j,s}(\eta_j, s) F_i$$

where $A_{j,s}(\eta_j, s) = a_i(\eta_j) s^q$, in which $q$ is a positive exponent signifying that larger individuals search a larger volume per unit time. The food intake is described using a type II functional response:

$$I_{j,s}\left(\eta_j, s, \mathrm{F}\right) = hs^m \frac{A_{j,s}\left(\eta_j, s\right) F_i}{c_{j,s}\left(\eta_j, s, \mathrm{F}\right) + hs^m}$$

Ingested food is assimilated with an efficiency $\epsilon_a$. Assimilated energy is first used to cover metabolic maintenance costs:

$$E(\eta_j, s, \mathrm{F}) = \epsilon_a I_{j,s}(\eta_j, s, \mathrm{F}) - ks^p$$

where $k$ is the size-specific constant of metabolic rate and $p$ is the size-scaling exponent of metabolic rate. Once metabolic maintenance costs are covered, a fraction $\psi$ of the available energy is used for reproduction and the remaining for somatic growth:

$$G_j(\eta_j, s, \mathrm{F}) = \begin{cases} E(\eta_j, s, \mathrm{F})(1 - \psi(s, s_m)) & E(\eta_j, s, \mathrm{F}) > 0 \\ 0 & \text{otherwise} \end{cases}$$

A smooth step function is used to describe how energy allocated to growth gradually changes from 1 to 0 around the body size at maturation $s_m$

$$\psi(s, s_m) = \left(1 + \left(\frac{s}{s_m}\right)^{-u}\right)^{-1} \left(\frac{s}{s_{\max}}\right)^{1-m}$$

In this equation, $s_{\max}$ is the maximum asymptotic body size, and $u$ determines the width of the transition for energy allocation from growth to reproduction. According to this expression, when the maximum asymptotic body size is reached, an individual allocates all its available energy to reproduction. An individual therefore produces offspring at a rate:

$$B(\eta_j, s, \mathrm{F}) = \begin{cases} \frac{\epsilon_b}{s_b} E(\eta_j, s, \mathrm{F}) \left(\psi(s, s_m)\right) & E(\eta_j, s, \mathrm{F}) > 0 \\ 0 & \text{otherwise} \end{cases}$$

where, $\epsilon_b$ is the efficiency at which assimilated energy is converted into offspring and $s_b$ is the size at birth.

Mortality decreases with body size following:

$$\delta(s) = \delta_p e^{-s}$$

Starvation mortality is not considered because at the ecological equilibrium, a population is viable (i.e. its density is positive) only if starvation conditions do not occur, i.e. if $E(\eta_j, s, \mathrm{F}) > 0$.

All assumptions discussed above pertain to the individual-level description. Without making further assumptions the population-level dynamics can be derived by bookkeeping following the Physiologically Structured Population Modeling approach[2]. The resulting ecological dynamics are:

$$\frac{dF_i}{dt} = \rho(F_{i\max} - F_i) - \sum_j \int_{s_b}^{\infty} h\, s^n \frac{A_{j,s}(\eta_j, s) F_i}{\sum_k A_{j,k}(\eta_j, s) F_k + h m^n} N_j(t, s) ds,$$

$$\frac{\partial N_j(t,s)}{\partial t} + \frac{\partial G_j(\eta_j, s, \mathrm{F}) N_j(t,s)}{\partial s} = -\delta(s) N_i(t, s),$$

and

$$G_j(\eta_j, s_b, \mathrm{F}) N_i(t, s_b) = \int_{s_b}^{\infty} B(\eta_j, s, \mathrm{F})\, N_j(t, s) ds$$

as the boundary condition for the population reproduction rate. Parameter values are determined from cross-species analysis of fish communities[1] and given in table SI6.1.

**Table SI6.1. Parameter values**

| Parameter | Description | Value | Unit |
|---|---|---|---|
| Diversifying lineage | | | |
| $s_m$ | Body size at maturation | evolving trait | g |
| $\eta$ | Feeding niche trait | evolving trait | - |
| $D$ | Distance between optima to feed on the resources | 1 | - |

| | | | |
|---|---|---|---|
| $\tau$ | Width of the Gaussian curve describing the degree of specialization to successfully attack a food resource | 1/3 | - |
| $\alpha$ | Maximum size-specific attack rate | 21.9 | $m^3$ $g^{-q}$ $d^{-1}$ |
| $h$ | Maximum food intake | 0.233 | $g^{1-m}$ $d^{-1}$ |
| $m$ | Exponent for maximum food intake | 0.75 | - |
| $q$ | Size-scaling exponent for search volume | 0.8 | - |
| $k$ | Standard metabolic rate | 0.0274 | $g^{1-p}$ $d^{-1}$ |
| $p$ | Size-scaling exponent of metabolic rate | 0.75 | - |
| $\epsilon_a$ | Assimilation efficiency | 0.6 | - |
| $u$ | Width of the transition for energy allocation from growth to reproduction | 10 | - |
| $s_{\max}$ | Maximum asymptotic body size | 1000 | - |
| $s_b$ | Body size at birth | 1 | g |
| $\epsilon_b$ | Efficiency of offspring production | 0.1 | - |
| $\delta_p$ | Size-dependent mortality rate | varied | $d^{-1}$ |
| Preexisting resources | | | |
| $\rho$ | Renewal rate of preexisting resources | 0.01 | $d^{-1}$ |
| $\phi$ | Total productivity of the habitat | varied | g $L^{-1}$ $d^{-1}$ |
| $n$ | Number of basal resources in the habitat | 2 | - |

Without making further assumptions the evolutionary dynamics can be obtained using standard methods of adaptive dynamics for size-structured populations[3]. These methods are incorporated into the PSPManalysis software package[4]. We use this package to determine whether diversification can occur or not for various levels of productivity and the body size at maturation, as well as the selection gradient of this life history trait (figure 4B).

***Model 3: Diversification and evolution of timing of diet shift***

We adopt a minimal model to describe a size-structured population with a diet shift[5]. The model describes the dynamics of a population of individuals using different food resources in consecutive stages of their life history. For simplicity, we assume that the feeding rate on a food resource that is used only early in life is constant, such that there is no density dependence at this life stage. This may occur when the difference in the productivity between the resources used in different life stages is very large, causing density dependence to regulate the population dynamics only in the stage feeding upon the resource with the lowest productivity. In our case, this resource is the resource used late in life.

We consider an ecomorph population $j$ to be composed of individuals characterized by their feeding niche trait $\eta_j$, body size $s$ and body size at diet shift $s_s$. Individuals are born with size $s_b$ and have access only to an early-life food resource, on which they feed at a rate $f_b$. They feed upon this resource until they reach body size $s_s$, when they shift their diet and can exploit a variety of resources $\mathrm{F} = (F_1 \dots F_n)$. Juvenile individuals mature and start to reproduce at a body size $s_m$. Individuals grow following:

$$G(\eta_j, s, \mathrm{F}) = \begin{cases} \epsilon_g f_b & \text{if } s < s_s \\ \epsilon_g \sum_{i=1}^{n} a_i(\eta_j) F_i & \text{otherwise} \end{cases}$$

and adults reproduce at a rate

$$B(\eta_j, s, \mathrm{F}) = \epsilon_b \sum_{i=1}^{n} a_i(\eta_j) F_i.$$

at a rate $g_1(s, \mathrm{F}) = \epsilon_g\, f_b$ before they reach size $s_s$, and a rate $g_2\,(\eta_j, s, \mathrm{F}) = \epsilon_g\, \sum_{i=1}^{n} a_i(\eta_j) F_i$ when they are larger. Adults reproduce at a rate $g_2\ = \epsilon_g\, \sum_{i=1}^{n} a_i(\eta_j) F_i$.

Mortality generally changes during life history. In particular, during an ontogenetic diet shift, organisms may experience such changes because a diet shift is often associated with a shift in habitat use and mortality may largely differ between habitats[6]. To reflect this, mortality is described as:

$$\delta_s(s) = \begin{cases} \delta_1 & \text{if } s < s_s \\ \delta_2 e^{-s} & \text{otherwise} \end{cases}$$

Using standard methods to formulate size-structured populations from individual life history processes[2] and after non-dimensionalizing the model following ref[5], we obtain the set of equations that described the ecological dynamics:

$$\frac{dF_i}{dt} = (\upsilon - \phi F_i) - \sum_j \int_{w_s}^{\infty} \gamma_2(\eta_j, w, \mathrm{F})\, M_{2,j}(t, w) dw,$$

$$\frac{\partial M_{1,j}(t, w)}{\partial t} + \frac{\partial \gamma_1(f_b) M_{1,j}(t, w)}{\partial w} = -\varsigma_1 M_{1,j}(t, w)$$

$$\frac{\partial M_{2,j}(t, w)}{\partial t} + \frac{\partial \gamma_2(\eta_j, w, \mathrm{F}) M_{2,j}(t, w)}{\partial w} = -\varsigma_2(w) M_{2,j}(t, w),$$

and the boundary condition for the diet shift

$$\gamma_2\,(\eta_j, w, \mathrm{F}) M_{2,j}(t, w_s) = \gamma_1(f_b) M_{1,j}(t, w_s)$$

and the population reproduction rate

$$\gamma_1(f_b) M_{1,j}(t, 0) = \beta\, \gamma_2\,(\eta_j, w, \mathrm{F}) \int_1^{\infty} M_2(t, w)\, dw.$$

In this system $\gamma_2\,(\eta_j, w, \mathrm{F}) =\ \sum_{i=1}^{n} a_i(\eta_j) F_i$ and $\varsigma_2(w) = \varsigma_c e^{-w}$. The scaled body size variable $w$ is related to the original body size measure following $w = (s - s_b)/(s_m - s_b)$ and maturation hence occurs at $w = 1$.

In the non-dimensionalized system the parameters relate to the parameters of the individual life history description presented above according to Table SI6.2. We adopt the parameter values in ref[5] for the life history description (see Table SI6.2), and the parameter values related to resource use after the diet shift are taken as in the model presented in the main text (see Table SI2.1).

**Table SI6.2. Non-dimensionalized parameter values**

| Description | Symbol | Relation with unscaled parameters | Value |
|---|---|---|---|
| Resource growth rate | $\phi$ | $\phi = \rho \sqrt{\frac{s_m - s_b}{\epsilon_g f_b}}$ | 0.01 |
| Mortality before the diet shift | $\varsigma_1$ | $\varsigma_1 = \delta_1 \sqrt{\frac{s_m - s_b}{\epsilon_g f_b}}$ | Varied (0.2, 0.25, 0.3) |
| Scaling factor of mortality after the diet shift | $\varsigma_c$ | $\varsigma_c = \delta_2 \sqrt{\frac{s_m - s_b}{\epsilon_g f_b}}$ | 0.5 |
| Adult fecundity scaled constant | $\beta$ | $\beta = \frac{(s_m - s_b)\epsilon_b}{\epsilon_g}$ | 0.01 |
| Body size at the diet shift | $w_s$ | $w_s = \frac{s_s - s_b}{s_m - s_b}$ | evolving trait |

We use the same suit of approaches as in model 2 to study the evolution of the feeding niche trait and the body size when individuals shift diet, which determines the timing of this shift. Therefore, the evolutionary dynamics can be obtained using standard methods of adaptive dynamics for size-structured populations[3]. We use the PSPManalysis software package[4] that implements these methods to determine whether diversification can occur or not for various levels of productivity and the body size at the diet shift, as well as the selection gradient of this life history trait (figure 4C).

**References**


1. Hartvig, M., Andersen, K. H. & Beyer, J. E. Food web framework for size-structured populations. *J. Theor. Biol.* **272**, 113–122 (2011).

2. de Roos, A. M. A Gentle Introduction to Physiologically Structured Population Models. in *Structured-Population Models in Marine, Terrestrial, and Freshwater Systems* (1997). doi:10.1007/978-1-4615-5973-3_5.

3. Durinx, M., Metz, J. A. J. & Meszéna, G. Adaptive dynamics for physiologically structured population models. *J. Math. Biol.* **56**, 673–742 (2008).

4. de Roos, A. M. PSPManalysis: Steady-state and bifurcation analysis of physiologically structured population models. *Methods Ecol. Evol.* **12**, 383–390 (2021).

5. Chaparro-Pedraza, P. C. & de Roos, A. M. Ecological changes with minor effect initiate evolution to delayed regime shifts. *Nat. Ecol. Evol.* **4**, 412–418 (2020).

6. Sánchez-Hernández, J., Nunn, A. D., Adams, C. E. & Amundsen, P. A. Causes and consequences of ontogenetic dietary shifts: a global synthesis using fish models. *Biol. Rev.* **94**, 539–554 (2019).

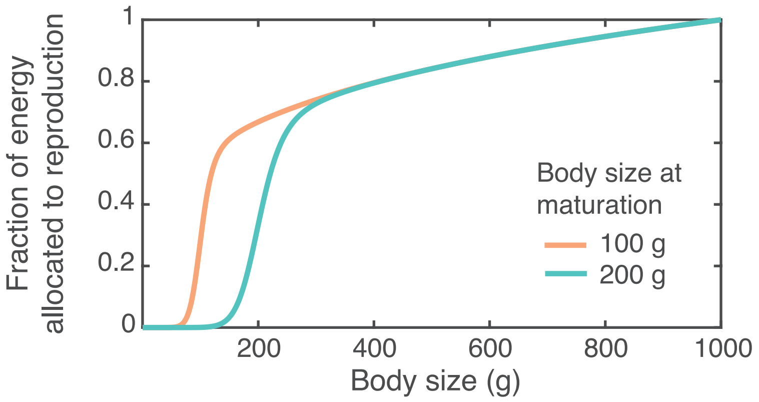


*Figure S1. Energy allocation and body size at maturation.*
Energy allocation into reproduction changes smoothly from 0 around the maturation size to 1 at the theoretical maximum asymptotic size (1000 g). Maturing at a small size enables an early onset of reproduction but reduces the growth rate. Because survival often increases with body size, the optimal body size at maturation depends on the tradeoff between reproductive investment and size-dependent survival.
